# From Attitudes to Actions: Bridging Conservation Planning Framework, Theory of Planned Behaviour and Social Values to Protect Charismatic Freshwater Fishes, the Mahseer

**DOI:** 10.64898/2025.12.26.696566

**Authors:** Prantik Das, V. V. Binoy

## Abstract

Mahseers are a group of iconic freshwater fishes native to the waterbodies of South, East, and South-East Asia, with significant ecological, socio-cultural, and livelihood value. The complexity of the socio-ecological contexts in which the mahseer conservation occurs demand an approach integrating behavioural dimensions of the individuals and groups directly or indirectly involved in it and social and cultural values they hold for these fishes with the conservation planning and implementation processes for better results. This study examined different facets of the mahseer conservation across three environmentally, geographically, and socio-culturally distinct Indian states - Karnataka, Assam, and Uttarakhand under a framework integrating Conservation Planning Framework (CPF) with the Theory of Planned Behaviour (TPB) and Social Values (SV). Across all the focal states, three key themes were constructed – ‘consensus between stakeholders,’ ‘communication and collaboration,’ and ‘values and moral responsibility.’ Our analyses revealed that while shared ecological concerns offered an opportunity for the collective conservation actions, inter-stakeholder conflicts, communication gaps, hesitation to collaborate, top-down governance, and restricted decision-making autonomy available to tribal and local communities acted as the barriers. Stakeholders from all three states demonstrated strong pro-conservation attitudes, moral responsibility, perceived capability to implement the conservation plans and culturally embedded value for mahseer, rooted in religious beliefs, tribal and local identities and recreational traditions. However, on ground analysis revealed a low ‘actual behavioural control’ amongst many stakeholders limiting translation of the positive elements present in their attitudes and values into tangible conservation outcomes. Our observations emphasise formalising community-based co-management, strengthening inter-departmental coordination, building conservation capacity, adopting socio-culturally grounded communication strategies and repositioning community fishing events as socio-ecological heritage to ensure stakeholder compliance of the existing mahseer conservation plans and improve its success. Furthermore, aligning conservation planning with behavioural drivers and social values could offer more vital insights for formulating inclusive strategies and policies for protecting mahseers and managing their habitats.

## 1. Introduction

India is home to 22 of the 56 mahseer species found across the globe (Fricke et al. 2025). These iconic, freshwater fishes (with some species exceeding 30 kg in size - megafishes), are taxonomically classified into three genera: *Tor*, *Neolissochilus* and *Naziritor* (Fricke et al. 2025; Froese and Pauly 2025). Amongst these, two species are critically endangered, two are endangered, one is vulnerable, three are near threatened, one is of least concern, five are data deficient and the rest continues as status not evaluated (Froese and Pauly 2025). The cultural diversity showcased by India can be seen reflecting also in the way in which mahseers are treated by the people in this nation. Evidence is available to show that mahseers enjoyed social significance from the time of the Indus Valley Civilisation (Hora 1956; Belcher 1998; Pinder et al. 2019). In Indian states of Karnataka, Maharashtra and parts of Uttarakhand, these fishes hold a sacred status and are often referred to as “God’s Fish” (Gupta et al. 2016; Pinder et al. 2019). Many communities from these regions consider killing and eating mahseers as inauspicious, and such activities may bring bad luck on those who do it (Dandekar 2011; Harad 2023). Similarly some tribal communities in the Northeastern states of India also revere these fishes (Dunbar 1915; Borah 2015; Sarma et al. 2018; Ruffner and Smith 2019). These beliefs can be seen translated into several community-protected “temple-based fish sanctuaries” to safeguard these revered fishes in Karnataka, Maharashtra and Uttarakhand (Gadgil 1991; Dandekar 2011; Katwate et al. 2014), along with several other government-local communities partnered and managed fish sanctuaries in Meghalaya (Sugnnan 1995; Rahman 2021). However, in North-Eastern states like Assam, Arunachal Pradesh, Nagaland and Sikkim, along with the Northern regions of West Bengal, local community members have been fishing mahseers for generations (Baruah and Sarma 2018) and these fishes often fetch high price in the market (Dunbar 1915; Sikkim Government Gazette 2021). They are also fished in other states such as Himachal Pradesh, Jammu and Kashmir, Odisha and Kerala (Gupta et al. 2020). Traditionally in India, mahseers have been angled for recreation (Nautiyal 2014; Gupta et al. 2015) and a 12^th^ Century Sanskrit text *Manasollasa* describes mahseers as valuable for both culinary and angling purposes (Hora 1953; Sadhale and Nene 2005). From the British colonial era anglers from different continents visited India to hook these mighty fishes (Nautiyal 2014; Gupta et al. 2015a). In recent times, catch and release recreational angling-based ecotourism boosted local economies by generating income, providing livelihood opportunities and employment for local communities (Everard and Kataria 2011; Pinder and Raghavan 2013) in Uttarakhand, Karnataka and Himachal Pradesh. Sadly, natural populations of many of these mahseer species are facing threats of extermination across India due to multiple pressures such as dam construction, siltation, introduction of alien species, community and mass fishing, illegal and destructive fishing, poaching, overfishing, deforestation and unregulated angling (Raghavan et al. 2011; Bhatt and Pandit 2016; Lewin et al. 2019).

Various plans are being implemented by both central and various state governments in India to conserve mahseers. Designating many species as state fishes (e.g. *T. putitora* by Jammu and Kashmir, Himachal Pradesh, Uttarakhand and Arunachal Pradesh; *T. mahanadicus* by Odisha; *T. tor* by Madhya Pradesh and *N. hexagonolepis* by Sikkim and Nagaland; Akhtar and Ciji 2023; Sikkim Government Gazette 2021), banning recreational angling in all protected areas across the nation (Ajay Dubey vs. NTCA: SLP No. 21339/2011; Gupta et al. 2015), captive breeding of many mahseers and restocking approximately 5,00,000 fingerlings into different rivers and lakes annually (Kulkarni and Ogale 1995; ICAR-DCFR 2016), etc. are few of such measures taken. Furthermore, the many state governments such as Assam, Bihar, Himachal Pradesh, Karnataka and Uttarakhand have implemented specific Fisheries Acts and Rules to prohibit fishing activity during the breeding season to protect migrating mahseer breeders (The Assam Fishery Rules, 1953; The Himachal Pradesh Fisheries Act, 1976; The Uttarakhand Fisheries Act, 2003 (Uttarakhand Act No. 2 of 2003)), lease specific stretches of rivers to NGOs as for conservation purposes, establish fish sanctuaries, protected areas, no fishing zones, ban of fishing and fish movement during closed seasons (The Karnataka Inland Fisheries (Conservation, Development and Regulation) Act, 1996 (Karnataka Act 27 of 2003); Bihar Jalkar Management (Ammendment) Act, 2018 (Bihar Act 13 of 2006)). Despite these efforts, mahseer populations across India continue to dwindle (Pinder et al. 2019; Sarma et al. 2022; Abass 2024). Evidences are abundant to support the argument: identification of all relevant individuals and groups affected by, or capable of influencing any step of the management plan (stakeholders; Decker et al. 1996; 2002) and fostering conservation-positive changes in their attitude and behaviour (Cinner 2018) is equally important to understanding ecological, breeding, genetics and taxonomic aspects of the focal species for conserving any wild population. Furthermore, excluding groups or individuals, especially those with whom the researchers and wildlife managers disagree (Marchini 2014) and the differences existing in the values attributed by different stakeholders to the wild species can significantly undermine conservation efforts (Marchini 2014), result in conservation policy failure, and may even lead to the human-human conflict (Dorow et al. 2009; Hunt et al. 2013; Arlinghaus et al. 2016). Although attempts to integrate human dimension into fish conservation are not novel (Cinner 2018), this source of knowledge remains underutilized both in policy making and on-ground-conservation actions to protect aquatic species in India. With the interest in mahseer conservation growing amongst the policymakers, government officials, scientists, anglers and local communities in India (Pinder et al. 2019; Roy and Sreenivasan 2025), it is vital to align perspectives, knowledge, attitudes, values, individual and collective ways of feeling and acting as well as trace out the socio-cultural, political and environmental factors determining human mahseer-relationship in different states of India to protect these apex predators in their natural habitats (Dacks et al. 2025).

One of the schemes popularly utilised for integrating human dimensions into wildlife conservation and management is the Conservation Planning Framework (CPF; Marchini et al. 2019). Studies based on CPF are conducted to find effective strategies for mitigating human-human conflicts and promoting human-wildlife coexistence (Pimid et al. 2022). CPF examines three fundamental aspects of conservation: (a) individual attributes (knowledge, attitudes, values), (b) social phenomena (governance, policies) and (c) social processes (decision-making, development; Bennett et al. 2017; Pimid et al. 2022). The CPF involves three essential stages, each encompassing multiple themes: situation assessment, decision-making and implementation, and monitoring and evaluation (Marchini et al. 2019; Zuluaga et al. 2020; detailed in Supplementary Materials SM 1). Since social values can guide what people perceive and the action chosen by them towards the focal organisms decisively (Chan et al. 2016) recent research strongly advised the integration of this concept into CPF (Pimid et al. 2022). Although value is a debatable topic due to its sensitivity to individual, context and culture, Chan et al. (2016) classifies the social values of humans towards nature and wildlife into three categories, viz., intrinsic (appreciation of wildlife and nature for its own sake; inherent values), instrumental (value of wildlife based on human material needs and usefulness; valuing for tangible benefits like livelihood) and relational (value of wildlife developed through cultivated relationships and meanings with humans that involves a sense of morality, responsibility, spiritual, identity, sense of belonging, personal and cultural connection with wildlife). Relational values built on the cultural beliefs and social norms prevalent in a community can in turn promote both intrinsic and instrumental values by influencing cultural and individual identity, moral responsibility to non-humans, social responsibility, environmental stewardship, etc. (Gupta et al. 2016; Thinley and Hartz-Karp 2019; Balmford et al. 2021). Hence the Intergovernmental Science-Policy Platform on Biodiversity and Ecosystem Service (IPBES) Value Assessment (2022) recommends utilising relational values for formulating conservation plans involving multi-stakeholders (IPBES 2022a, 2022b; Pascual et al. 2023).

Along with the knowledge of different dimensions of the social values, insights on the psychological underpinnings of the responses towards a conservation issue by different stakeholders are also essential to promote sustainable pro-conservation behaviours and the environmental stewardship in a society by bringing positive behaviour change. In many contexts, successful induction of the desired changes in the behaviours of the targeted stakeholders can even reduce the cost of expensive interventions and hence catalyse the momentum of the conservation actions often afflicted by the limited resource availability (Echols et al. 2019). One of the robust and widely used socio-psychological models of human behaviour in conservation scenarios is the Theory of Planned Behaviour (TPB; Ajzen 1985, 1991). According to TPB the strongest predictor of what a person or a group does in a given context is their intention (Zhong et al. 2019). However three factors popularly known as the motivational factors: attitudes (Ullah et al. 2021), societal norms and pressures, and perceived control over one’s own behaviour (Ajzen 1985, 1991) - have profound influence on the intention and hence the choices they make in a decision making context (Sánchez et al. 2018). The first component, attitude, plays a central role in TPB, which significantly shapes how an individual perceives and evaluates a behaviour associated with a particular organism. This evaluative judgment is largely influenced by the individual’s underlying beliefs and the context in which the behaviour occurs (Ullah et al. 2021). Social norms or social pressures originate from the perceived expectations of the significant others (family, peers, or community leaders) and the actors’ belief about whether these important people would support/disapprove their chosen action (Liang et al. 2018). Meanwhile, the perceived behavioural control (PBC) is related to the self-efficacy and the belief of the ease/difficulty of performing the behaviour (Sánchez et al. 2018; Empidi and Emang 2021; SM 2). Therefore, positive attitudes, supportive social norms and strong perceived control could encourage positive changes in the behaviours (Farani et al. 2021; Karimi and Ataei 2022; Batool et al. 2024). In addition to these three motivating factors, other non-motivating external factors like availability of time, resources, facilitating conditions, autonomy in decision making, etc. that can also directly influence the behaviour as well as the PBC, are known as actual behavioural control (ABC; Ajzen 1991, 2020).

CPF offers a structured and stepwise approach for effective conservation planning, decision-making and conflict management (Marchini et al. 2019; Pimid et al. 2022), while explaining how individual behavioural intentions are determined by the attitudes, subjective norms and PBC is the aim of the TPB (Ajzen 1985, 1991). Another important aspect of the conservation behaviour at individual and community level, is the attribution of values, to various components of nature including non-human life forms, and its role in directing people’s action for conservation is elaborated by the widely studied social values approach (Chan et al. 2016; Balmford et al. 2021; Pascual et al. 2023). These help reveal a deeper normative foundation as to why people care about conservation in the first place, based on how they value wildlife. These values influence TPB variables such as attitudes and norms and, through them; they also influence the CPF outcomes. For example, strong relational values may increase willingness to engage in conservation, while a primarily instrumental outlook might lead to resistance if conservation is seen as economically damaging (IPBES 2022a; 2022b; Jolly and Stronza 2025; Iqbal et al. 2025).

Although these three models may look independent, a closer examination reveals overlap between many concepts on which they are built on, hence offer a great potential for integration thereby enhancing the effectiveness of conservation planning (Fig. 1). Conservation behaviour being complex in nature, an approach amalgamating different frameworks utilised for explaining it could enhance the efficacy of interventions and policies designed keeping such ideas in the focus. For instance, the situation assessment phase of the CPF is dedicated for understanding knowledge, perceptions, and existing plans of the stakeholders, which aligns closely with the two major components of TPB: attitudes and subjective norms. Hence, incorporating the attitude component of TPB into the situation assessment could help better understand views and pro-conservation behaviours of stakeholders at the individual level thereby enabling policy makers to set clear goals and more accurate identification of the priority areas for conservation. It is a well-known fact that social values, more specifically the intrinsic, instrumental as well the relational values have profound effects on the opinions and actions of the stakeholders and also the motivational factors deciding them (Brackhane et al. 2019; Jolly and Stronza 2025). Intrinsic and relational values shape stakeholders’ attitudes and perceptions of control over their actions (PBC) thus their decision-making 9 (Dhee et al. 2019; Jolly et al. 2022), while knowledge of the instrumental values can help in managing conservation barriers. For instance, whether individuals perceiving conservation efforts as resource-intensive and lacking or having tangible or intangible benefits, could decide their motivation to act. Including these concepts of values could help in tightly bridging the first and second (decision-making and implementation) phases of CPF. Aligning individual interests, expectations, and normative practices with the rules and regulations promoted by institutions and the broader community, which these individuals are a part of, can help reduce the likelihood of conflict (Reed et al. 2014). Furthermore, CPF tends to take each stakeholder group as a collective with common interests and often fail to capture individual-level variations present in the mindset and behaviours of the members while considering decision-making, governance, communication and ensuring the involvement of diverse community members (second phase of CPF). However, TPB offers a more granular understanding of these aspects. Additionally PBC and ABC components of TPB can elucidate the “conservation barriers” discussed by the CPF by throwing light on how individuals’ perceived and actual control over action can impact outcomes of the conservation plans.

**Fig. 1.**
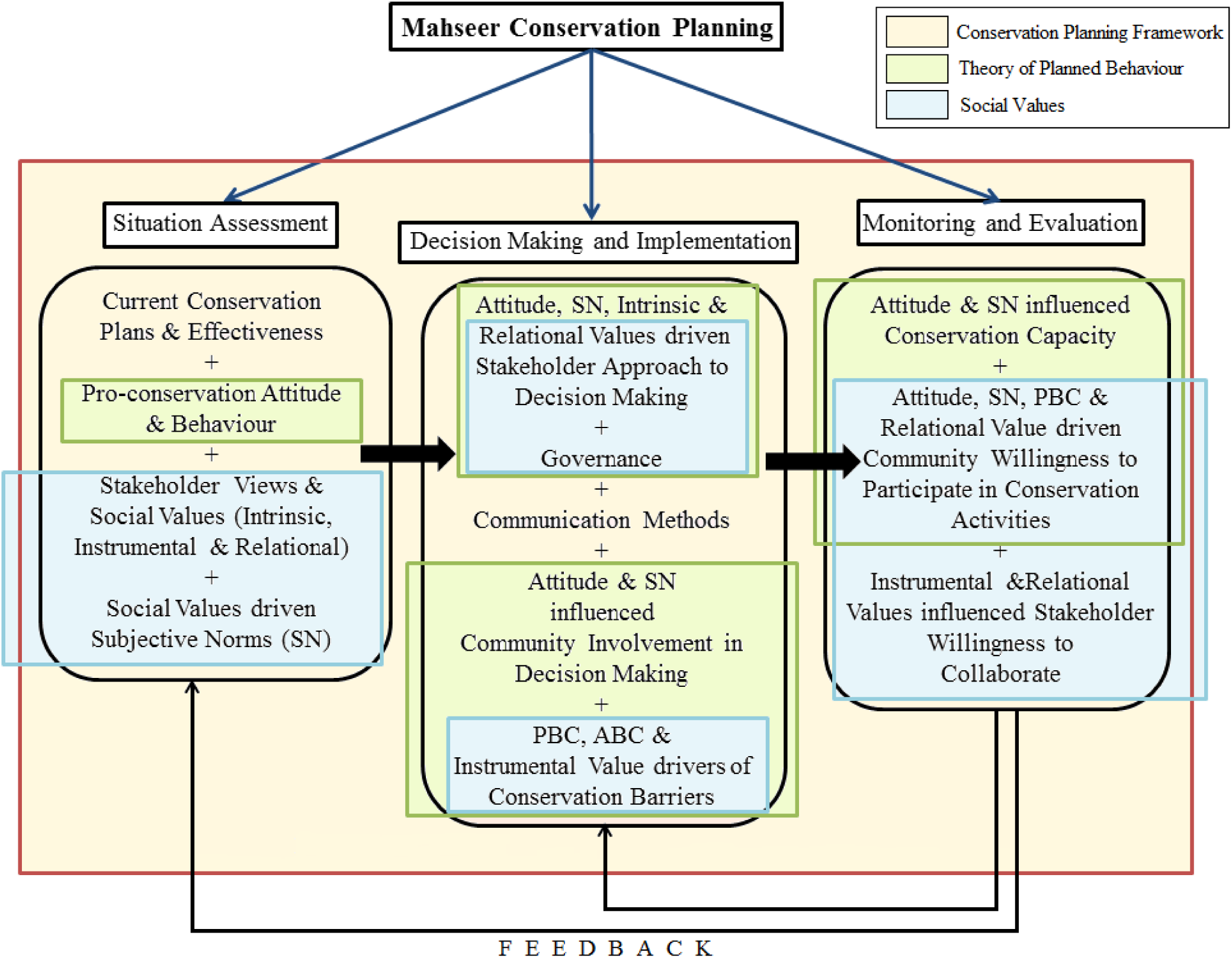
Integration of the Theory of Planned Behaviour (TPB) and Social Values into the Conservation Planning Framework (CPF). (SN: Subjective Norms; PBC: Perceived Behavioural Control; ABC: Actual Behavioural Control).

Monitoring and evaluation, the third phase of CPF, can also be benefited from the incorporation of TPB and social values. Development of a communication and governance plan considering the attitudes, norms and the determinants of the behavioural control of both individuals and groups of stakeholders can be useful in translating their conservation capacity into action and willingness to work for it. The impact of such an approach will be synergised when relational and instrumental values, with the potential to explain how stakeholders’ sense of morality, responsibility, belongingness, identity, personal and cultural relationships with wildlife and nature transform into the willingness to collaborate and engage in conservation-related activities, is also kept simultaneously in the focus.

Hence an integrated model where CPF is used as the structural backbone to outline the “what” and “how” of the conservation by groups, TPB as the individualistic behavioural mechanism explaining “why” some individuals act or while others fail to do so, and social values as the drivers of socio-cultural underpinnings; the “deeper why” behind behaviours and decisions at the levels of individuals and collective could provide a more comprehensive, stakeholder-oriented and sustainable foundation for conservation planning and decision making. Furthermore, including a feedback loop from the third phase of TPB and value integrated CPF to the first and second, will be beneficial in strengthening this framework by the timely incorporation of the new and constantly evolving dimensions of the attitudes, norms, insights and other determinants of actions (Marchini et al. 2019; Zuluaga et al. 2020; Pimid et al. 2022). Even though numerous stand-alone CPF and TPB based studies have been conducted (Marchini et al. 2019; Pimid et al. 2022) to explain, predict and manage a range of environmental and conservation issues (Renzi and Klobas 2008; Oztekin et al. 2017; Hill et al. 2019; Liu et al. 2021; Nageotte and Buck 2023; Przymuszala 2023; 2024; Savari and Khaleghi 2023), and in the recent past many researchers have begun incorporating social values into CPF (Pimid et al. 2022) and environmental planning (Vatn et al. 2024), up to our knowledge, any attempt to integrate these three important components for wildlife conservation has not been undertaken till date.

Considering the cultural, environmental and stakeholder diversity this novel approach integrating values, TPB and CPF can improve management and conservation of the much-important yet attention-deficient freshwater fish species such as the mahseers and their habitats in India where conservation discourses and programmes are traditionally focused on ecological and biological dimensions and often overlooks the socio-psychological aspects of human behaviour (Pinder et al. 2019; Das and Binoy 2024, 2025a, 2025b). Building on the integrated framework, the current study aims to address the following key questions:

1. Can the Conservation Planning Framework (CPF), Theory of Planned Behaviour (TPB), and Social Values be integrated into a single framework to examine multi-stakeholder attitudes, perceptions, values, and behaviours oriented towards the conservation of mahseer?
2. How does such an integrated framework help in understanding the points of agreement and variance in the mahseer conservation intentions and behaviours of the stakeholders of culturally distinct social-ecosystems?

## 2. Methodology

### 2.1. Study Areas

The present study focused on different stakeholders from three Indian states viz. Karnataka, Uttarakhand, and Assam are well known for the presence of mahseers. However, we had to restrict our analysis to five species: *Tor khudree* (Deccan mahseer), *T. putitora* (golden mahseer), *T. remadevii* (humpback/orange-finned mahseer), *T. mosal* (mosal mahseer) and *Neolissochilus hexagonolepis* (chocolate mahseer), due to the existing ambiguity over taxonomy, distribution and insufficient data availability (Data Deficient) for *T. tor* (deep-bodied mahseer; Rayamajhi et al. 2018) and *T. dukai* (Devi and Boguskaya 2009) which are also distributed in these regions. Key stakeholders were selected for the interviews and FGDs employing purposive sampling followed by snowball sampling methods. Multiple districts across these three states were selected to conduct the interviews and FGDs of the stakeholders are provided in detail in the following sections:

#### (a) Karnataka

The study in Karnataka was conducted in four districts located within the Cauvery River basin: Kodagu (Coorg), Hassan, Mandya and Ramanagara (Fig. 2). These districts are recognised biodiversity hotspots and habitat for both *T. khudree* and *T. remadevii* (Pinder et al. 2019). The former is found in rivers of central and southern India, (Ogale 2002; Pinder et al. 2019), and the latter is endemic to the Cauvery River basin in Karnataka, Kerala and Tamil Nadu (Kurup and Radhakrishnan 2007; 2011). Kodagu hosts the only state-run mahseer hatchery (Harangi hatchery; producing *T. khudree* fingerlings), and is a primary ranching site for *T. khudree* in the Cauvery. Interestingly, the first release of the mahseer fingerlings in this state by Tata Power’s Lonavala hatchery in the early 1990s, were in the Kodagu stretch of Cauvery (Kulkarni and Ogale 1995). Conservation actors such as the Chendanda Clan of the Kodava community, and two major recreational angling associations of the Karnataka state, WASI and Coorg Wildlife Society (CWS) have been active in this region since the 1970s. Mahseers holds a high religious and culture reverence in this are and are called ‘*devaru meenu*’ (God’s fish) in the study areas except in Kodagu and Mandya districts (here vernacular name is ‘*bili*’: white or ‘*gande meenu*’: back fish). Hence, the state Fisheries Department-declared temple-based fish sanctuary (*matsyadhamas*) for mahseers in Ramanathapura in Hassan district (Gokhale and Chandran 2009) along with multiple other formal, non-temple based fish sanctuaries in places like Kodagu (Harangi Dam to Kudige Bridge, Valnoor Cauvery stretch) and Mandya (Shivanasamudra Fish Sanctuary), account for a total of 22 fish sanctuaries in the state (Karnataka Handbook of Fisheries Statistics 2020; Roy and Sreenivasan 2025). In Mandya and Ramanagara districts, the Cauvery River flows through a Protected Forest Area (PA) that has been declared as the Cauvery Wildlife Sanctuary (CWLS) known for harbouring pristine and untouched wild populations of the two mahseer species (Roy and Sreenivasan 2025). A total of 23 interviews of the selected individual stakeholder and one FGD (n=19) were conducted in the English, Hindi and Kannada languages (Table 1). The anglers from the Wildlife Association of South India (WASI) operating mainly in Mandya and Ramanagara districts, participated in the FGD conducted in Bangalore Urban district.

**Fig. 2.**
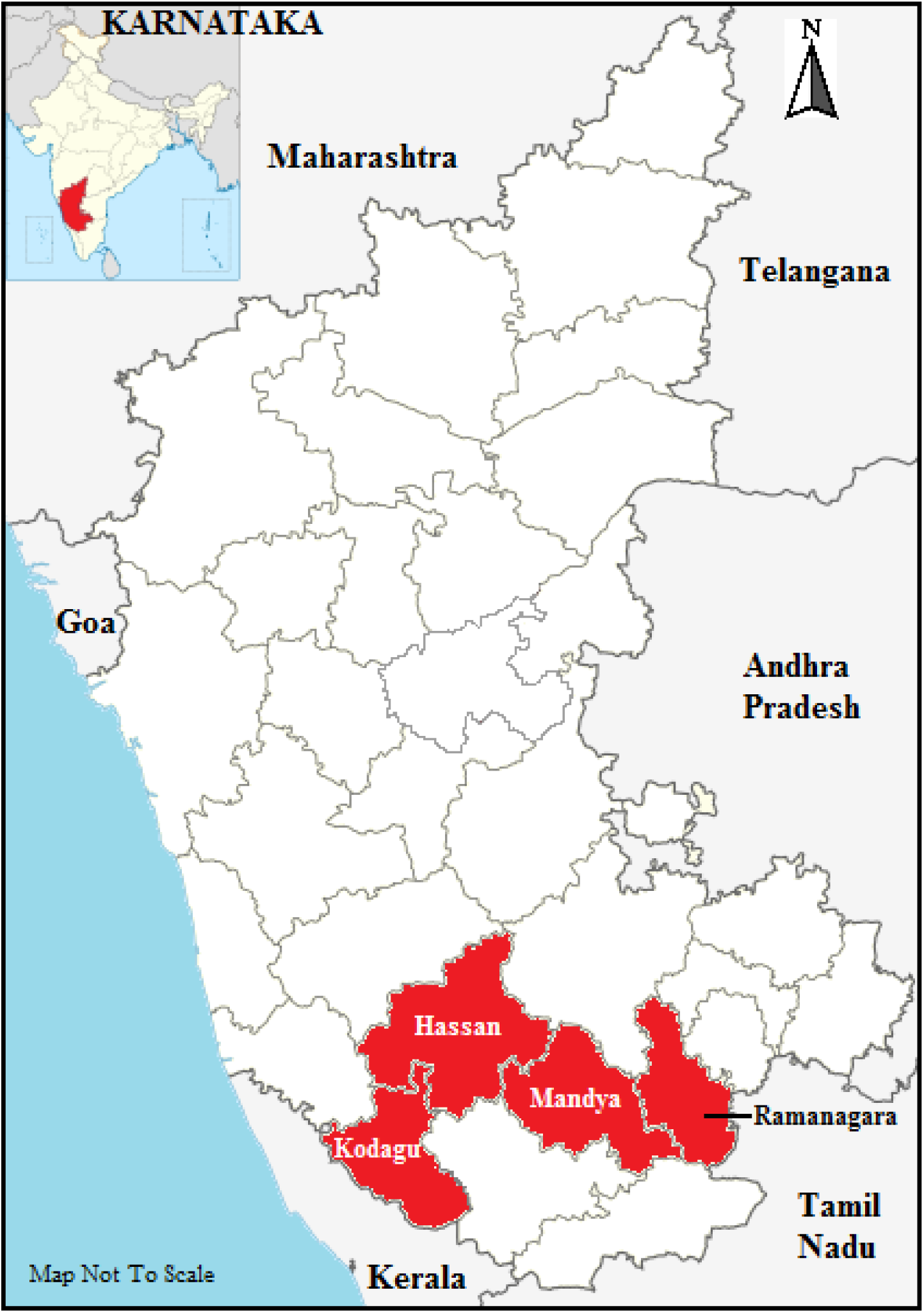
The districts in the Indian state of Karnataka where the study was carried out. Map not to scale.

**Table 1.** List of stakeholders from three Indian states interviewed and participated in the Focused Group Discussion (FGD) conducted for the study.

| No. | States | Species Focused | Stakeholders | Number of Participants |
| --- | --- | --- | --- | --- |
| 1. | Karnataka | *T. khudree*<br>*T. remadevii* | Fisheries Department | 3 |
|  |  |  | Forest Department | 5 |
|  |  |  | Recreational Anglers Associations | 6 |
|  |  |  |  | 19 (FGD) |
|  |  |  | Research and Educational Colleges/Institutes/Universities | 3 |
|  |  |  | Temple employed vendors at bank/locals (Ramanathapura Temple Fish-based Fish Sanctuary) | 6 |
| 2. | Assam | *T. putitora*<br>*N. hexagonolepis* | Fisheries Department | 5 |
|  |  |  | Forest Department | 4 |
|  |  |  |  | 11 (FGD) |
|  |  |  | Research and Educational Colleges/Institutes/Universities | 5 |
|  |  |  | Recreational Anglers Associations | 12 (FGD) |
|  |  |  | Pond and Lake/*Pukhuri* Managers | 6 |
|  |  |  | Local NGOs<br>(Aqua-tourism Centre-cum-Hatchery) | 2 |
|  |  |  | Local Community (Residents of Nameri National Park Fringe Village/Buffer Area) | 16 |
|  |  |  | Traditional Community Fishing Participants<br>(Members of Tribal Communities - Tiwa, Karbi, Bodo, and Rabha tribes, and Non-tribal Assamese Communities) | 40 |
|  |  |  | Community Fishing Festival/*Mela* Organising Committee<br>(Also includes members of Karbi and Gobha-Tiwa *Raja* 's/King's Court) | 11 |
|  |  |  | Politically Elected Members of Tiwa (Lalung) Autonomous Council | 2 |
| 3. | Uttarakhand | *T. putitora* | Fisheries Department | 8 |
|  |  |  | Forest Department | 8 |
|  |  |  | Research and Educational Colleges/Institutes/Universities | 4 |
|  |  |  | Recreational Angling Company<br>(Anglers-cum-Guides) | 7 |
|  |  |  | Local NGOs<br>(Eco-tourism, Research, Biodiversity<br>Conservation and Societal Research-based<br>organisations) | 5 |
|  |  |  | Local Fishermen-cum-Participants of<br>Community Fishing Events | 12 |

#### (b) Assam

The study in Assam was undertaken in four districts: Kamrup (Metropolitan), Morigaon, Nagaon and Sonitpur (Fig. 3) known for the presence of both *T. putitora* (golden or Himalayan mahseer) known locally as ‘*sonali pithia*’, and *Neolissochilus hexagonolepis* (chocolate mahseer; ‘*boka pithia*’). Assam is well known for the Brahmaputra River system, and the tributaries of this river flowing through the Nameri National Park (NP) in Tezpur, Sonitpur district, supports abundant wild mahseer populations (Borgohain 2015). Assam is also the only state in India with two ICAR-DCFR funded mahseer hatcheries; one within the Eco-Camp of NP and the second in Nagaon district (Borgohain 2015; Das and Gogoi 2015). The NP hatchery is operated in collaboration with the Indian Council of Agricultural Research - Directorate of Coldwater Fisheries Research (ICAR-DCFR) and the Assam Bhorelli Angling and Conservation Association (ABACA). In Nagaon, *T. putitora* is bred with the support from ICAR-DCFR and this facility hosts a unique aqua-tourism centre on the bank of the Mahrul *beel* (wetlands/lakes). This centre is also known for hosting annual National Angling Competitions and Fish Festivals attracting both anglers and the general public (Das et al. 2023). These initiatives have also contributed immensely to the local community by generating employment opportunities (Das et al. 2023). Moreover, the ICAR-DCFR has undertaken initiatives to ranch mahseers in various lakes across Guwahati, Kamrup (M) and Tezpur, Sonitpur and launched “Mahseer Watching” programmes along the lines of bird watching (Baruah 2018; Baruah and Sarma 2018) to increase public awareness of these fishes.

**Fig. 3.**
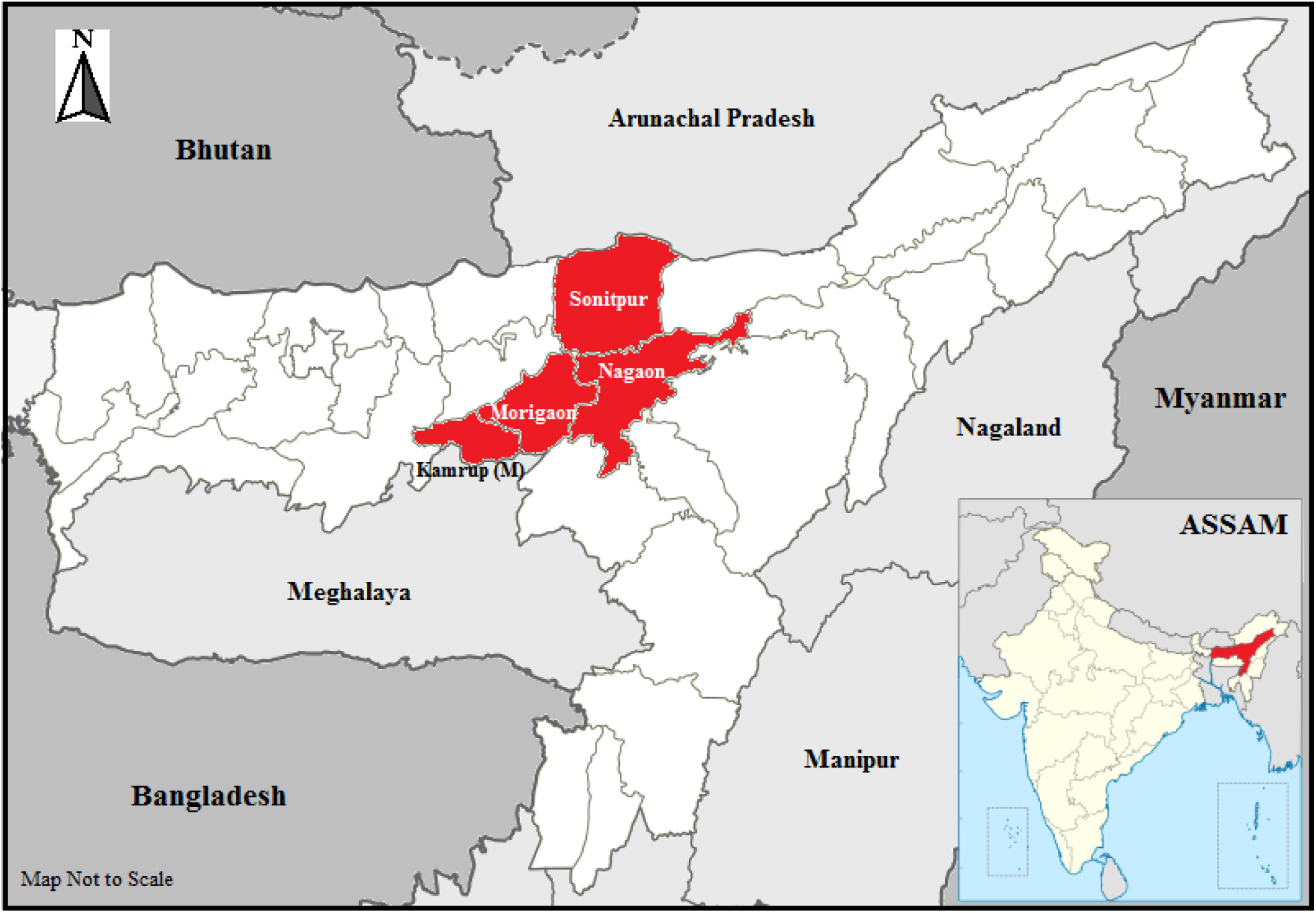
The districts in the Indian state of Assam where the study was carried out. Map not to scale.

Assam is also famous for traditional community fishing festivals. During the *Magh Bihu ‘Uruka’*, the harvest festival, multiple Tribal communities of this state such as Tiwa, Karbi, Bodo, along with the non-tribal Assamese people participate in such events to celebrate their identity, heritage and culture (Baruah 2017). For instance, Baghjap village in Mayong block (Morigaon district) conducts a three-day *Junbeel Mela*, annually. Community fishing, with participants from the Tiwa, Karbi, Bodo, Khasi and Jaintia tribes (Doloi 2024), is one of the main attractions of this cultural festival initiated by the Tiwa king with the help of Ahom kings of Assam and Jaintia kings of Meghalaya in 14^th^ or 15^th^ century (Roy 2017). Similarly in Kamrup (M), Maloibari village in Dimoria block hosts a historic fishing festival started in the early 19th century by the Kings of Tentelia (Telelia) and Dimoria (Das and Das 2020). Here, people irrespective of their social and religious backgrounds enjoy fishing in three *beel* namely Parokhali, Bomani and Jalikhora connected to the Digaru (tributary) and Kolong rivers (distributaries of Brahmaputra River). These festivals attract tourists and media from all over the country. However, the commercialisation and uncontrolled use of non-traditional equipment for fishing has negatively impacted the mahseers and other ichthyo- and avi-fauna of these aquatic ecosystems (Deka and Sharma 2013; Kalita et al. 2016; Medhi and Sharma 2017; Dutta et al. 2023). In Assam 91 interviews and two FGDs (number of participants = 11 and 12) were conducted in Assamese, English and Hindi languages (Table 1).

#### (c) Uttarakhand

Historical records indicate that in Uttarakhand *T. putitora* was introduced into Bhimtal and Nainital lakes in Nainital district in 1858 by British official Sir Henry Ramsay for the purpose of recreational angling (Walker 1888). In 2001, Uttarakhand recognised this species as their State Fish. Nainital district hosts the only central-government run *T. putitora* hatchery in India, operated jointly by the ICAR-DCFR and Nainital District Fisheries Department. Ramganga and Kosi rivers flowing through Corbett National Park and Kalagarh Tiger Reserve present in this district (Johal et al. 1994; Johnsingh et al. 2006) also harbours populations of mahseer. Furthermore, the Pancheshwar region of Champawat district, situated at the confluence of the Saryu and Mahakali Rivers near the India-Nepal border, is a globally acclaimed premier destination for *T. putitora* recreational angling with several private angling companies such as The Himalayan Outback, Pancheshwar Fishing, etc. operating here. The key spawning grounds for *T. putitora* have been identified across multiple rivers: Song and Tons Rivers (tributaries of the rivers Yamuna and Ganga) in Dehradun, Aglar in Tehri Garhwal, Nayar and Khoh in Pauri Garhwal (Bhatt et al. 2004); and the Mandal River in Nainital (Atkore et al. 2011).

Uttarakhand is also famous for *Maund* (or *Maun*) *Matsya Mela*, a century-old annual community fishing festival conducted in the Aglar River (tributary of Yamuna River), which unfortunately coincides with the onset of monsoon and fish breeding season (the months of June and July; Sharma et al. 2016). Started by the King of Tehri, in the mid-1860s, community fishing, primarily undertaken by the members of the Jaunpuri community from different villages from the Jaunpur areas of Tehri Garhwal and Dehradun districts (Sundriyal and Kumar 2019) is the main component of this festival attracting tourists and media from all over the country. Sadly, the use of a toxin powder, made from the bark, seeds and leaves of the Timur plant (*Zanthoxylum armatum*), to paralyse fishes in the river during the festival (Sharma et al. 2016; Singh et al. 2016), adversely affecting mahseers and other fish populations have been reported by many authors (Sharma et al. 2016; Sundriyal and Kumar 2019; Uniyal and Uniyal 2021). Stakeholders (n = 44) from the study area comprising Dehradun, Tehri Garhwal, Nainital and Champawat districts (Fig. 4) were interviewed in Hindi language. These districts are also known for several non-formal temple pools and fish sanctuaries such as along the Garjiya Devi temple, the Kosi River stretch at Khairna, and *Nal-Damyanti Tal* in Bhimtal in Nainital district which support healthy wild populations of mahseers (Dandekar 2013; Kattyayani 2016; Baruah et al. 2022).

**Fig. 4.**
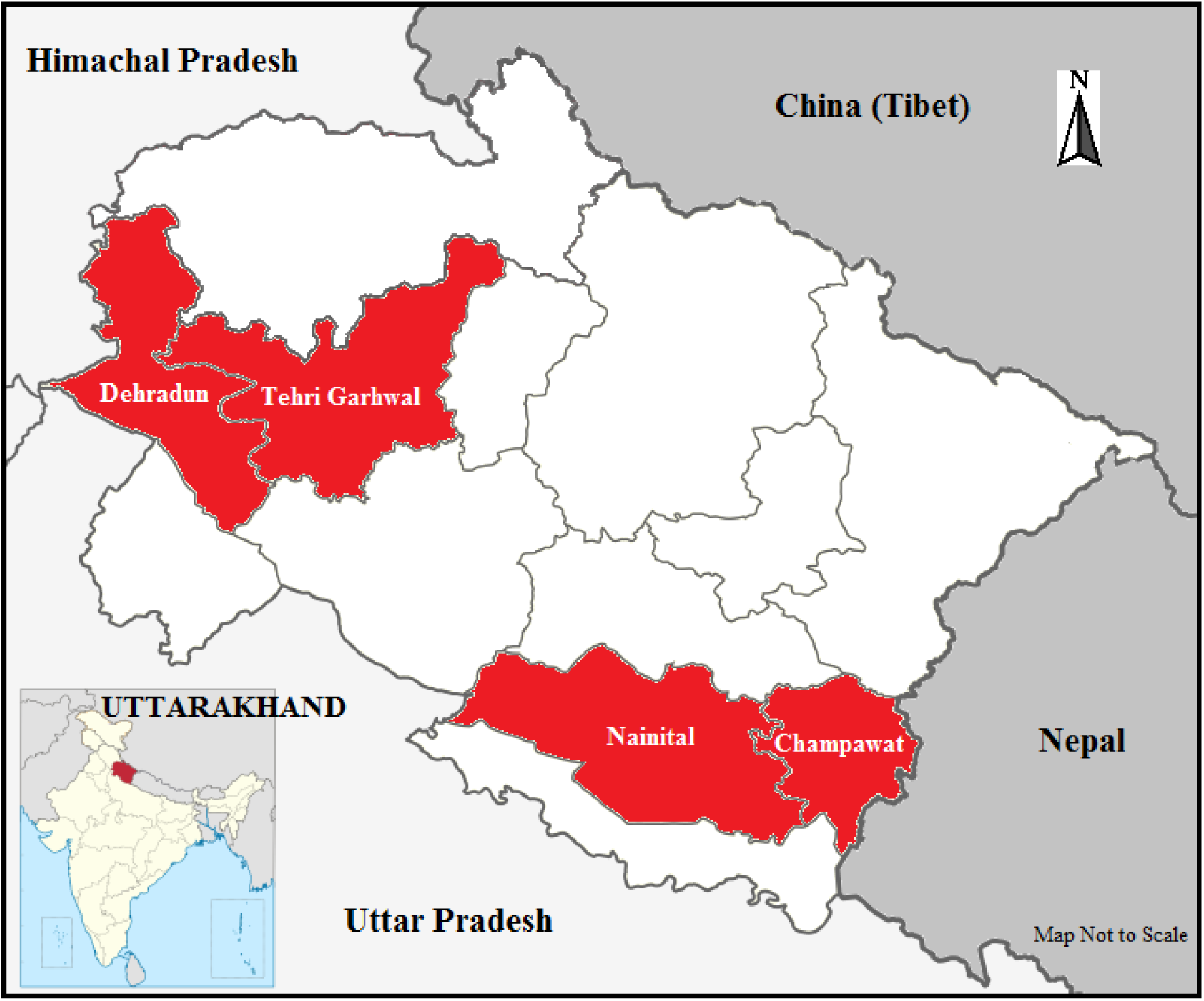
The districts in the Indian state of Uttarakhand where the study was carried out. Map not to scale.

### 2.2. Data Collection

In-depth, semi-structured, discursive, one-on-one interviews of the stakeholders were conducted following a list of questions reflecting multiple aspects of CPF and TPB (SM 3) which lasted between 15-45 minutes. In addition to these individual interviews, a total of three focus group discussions (FGDs; 1 in Karnataka (n = 19) and 2 in Assam (n = 11, 12) were conducted with the participants who preferred group discussions over interviews. We ensured that all interviews and FGDs adhered to the required institutional ethical guidelines for conducting non-invasive human research. Interviews and FGD were audio recorded, written notes taken and photographs captured after acquiring oral or written informed consent from the participants. The participants were assured of the anonymity of their names, personal details and designations to protect their identity and privacy.

### 2.3. Data Analysis

The English interviews were transcribed verbatim using Otter.ai and Rev.ai while the others were transcribed manually into English. The transcripts were then manually cross-checked for inconsistencies by referring to the audio recordings. Additionally, the field notes were reviewed for additional contextual support (Nkansah-Dwamena 2023). These transcripts were subjected to the thematic Qualitative Data Analysis (QDA) following a hybrid deductive-inductive coding approach to ensure greater qualitative depth and rigor (Fereday and Muir-Chochrane 2006; Braun and Clarke 2013; 2019; 2021; Byrne 2022; Proudfoot et al. 2022). The thematic QDA followed six steps, adapted from Braun and Clarke (2006) involving transcribing, data familiarisation, initial code generation (deductive codes based on CPF and TPB and additional inductive codes), organising codes into meaningful themes, reviewing and refining themes, and defining themes. As specified by Braun and Clarke (2021), thematic QDA is often reflexive in nature (Reflexive Thematic Analysis, RTA; Terry et al. 2017; Braun and Clarke 2019; Braun et al. 2022). The RTA focuses on the researcher’s knowledge, reflexivity, subjectivity, contextual specificity as well as on transparency, thoughtful and interpretative depth of the data. Differing from the conventional coding analysis, RTA rooted in constructivist interpretive approach, considers ‘replicability’ and inter-coder reliability unsuitable and inappropriate (Braun and Clarke 2019; 2021). Hence the entire data set was coded by a single coder (PD) well versed with the methodologies of RTA.

## 3. Results

### Karnataka

The situation assessment (CPF) revealed river conservation leases to angler and NGOs (frequency of codes from interviews and FGDs, Fr = 36; Fig. 5), issuing formal angling licenses (36), declaration of fish sanctuaries (42), conservation breeding of *T. khudree* (36), angling training (25), and anti-poaching camps (5) as the active intervention plans existing for the protection of mahseer populations in Karnataka. The stakeholder attitudes (TPB) were generally positive (36) in nature and many were of the opinion that these activities are useful in increasing the numbers of the wild mahseers and reducing illegal and destructive fishing activities.

**Fig. 5.**
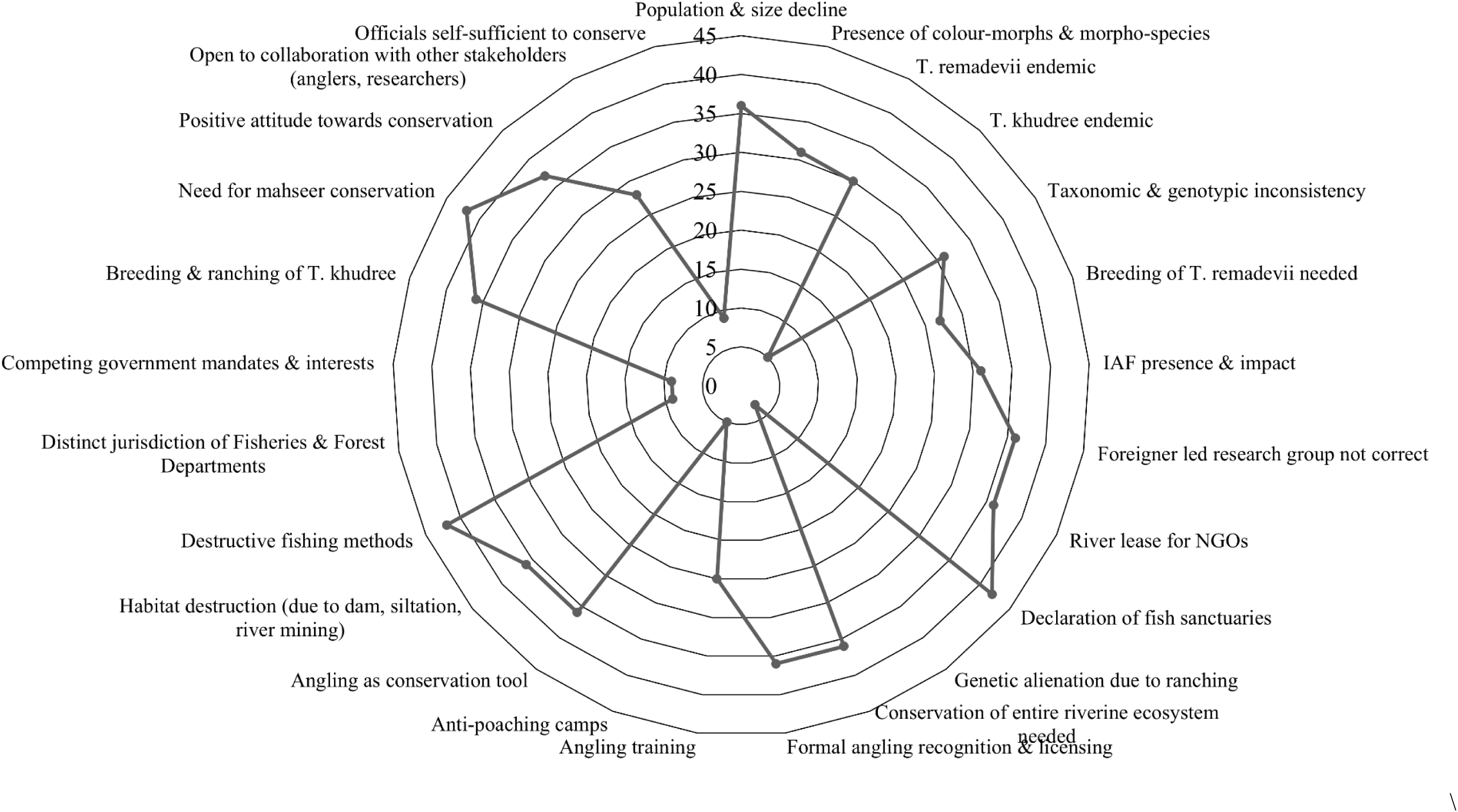
Radar chart depicting the frequency of codes used for generating the theme ‘consensus between stakeholder’ from the reflexive thematic analysis conducted on the interviews and FGDs from Karnataka.

‘Consensus between stakeholders’ was an important theme generated from the Karnataka data. This theme appeared in multiple contexts ranging from the taxonomic ambiguity and research leadership to the jurisdiction of the conservation activities. Many stakeholder groups were aware of and subscribed to the ongoing debates on the taxonomy of mahseers, and due to the presence of colour morphs and morpho-species they believed that rivers of Karnataka may be harbouring many more species of mahseer (Fr = 31; Fig. 5) than the currently known. This disagreement was found resonating also in the case of endemic status attributed to *T. remadevii* (30) and *T. khudree* (5).

> Quote 1: *“I think that T. remadevii and T. khudree are both endemic to the Cauvery River in Karnataka”. Quote 2: “Only T. remadevii is endemic to the Cauvery River in Karnataka, T. khudree is endemic to the rivers in Maharashtra and were introduced here”.*

The perception reflected in quote 1 was more prominent amongst the staff from the fisheries department (5) while quote 2 the anglers (25) and the forest department staff (5). However, respondents (36) from multiple stakeholder groups shared their dissatisfaction over ‘foreign scientist’ led mahseer research, especially in the area of taxonomy. In their opinion short-term research programmes run by the researchers from abroad fail to elucidate a holistic picture of the local mahseer populations and hence a dedicated long term plan lead by Indian-scientists is a need of the hour.

> Quote 3: *“We have our own jurisdiction, the forest department has their own. Aquatic organisms come under our purview while the terrestrial and arboreal animals are under them”.*

This statement highlights the lack of consensus among government departments involved in conserving natural mahseer populations. Although mahseer habitats lie within the rivers that traverse protected forest areas, these fish lack legal protection as they are not enlisted under any schedule of the Wildlife Protection Act (WPA) 1972. While the fisheries department breeds juvenile mahseers, the release sites may fall within protected forest areas, creating jurisdictional overlaps and disputes (9). Even though the stakeholder attitudes were generally positive, their willingness to be involved in conservation activities varied. The anglers and researchers demonstrated their openness to collaborate with other stakeholders (28), while the staff of the governmental departments were hesitant to collaborate, displaying a strong sense of self-efficacy (9). These jurisdictional tensions and conflicts arise from mismatched mandates (9), conflicting interests and underlying causes of political and power dynamics, which can often slow, hinder or derail coordinated wildlife conservation efforts (Lele et al. 2010; Jacobs et al. 2020; Riechers et al. 2025).

‘Communication and collaboration’ was another theme developed that closely aligned with the decision-making and implementation, the second component of CPF as well as the subjective norms and perceived control over behaviour (TPB). The sub-themes that came out under this category, communication gaps (42; Fig. 6), lack of collaborative mechanisms (34), and interdepartmental divergences and frictions (9) revealed more than technical shortcomings that could be patched easily in the existing mechanism for promoting mahseer conservation. Absence of credible communication channels (6) could further intensify the misalignment between stakeholders and delay the translation of positive attitudes present into the expected actions. Although social media was recognised as a tool for awareness creation (25) many identified this medium also as a promoter of unsustainable behaviours (25). Therefore, establishing a platform where representatives from the society and stakeholders can be involved in bidirectional communication (Bickford et al. 2012) can help in possibly resolving the clash of interests to a certain extent through the reconciliation of differences. Such a platform and active stakeholder interaction promoted by it, has the potential to bring about reconciliation by inculcating a sense of duty and moral responsibility towards the mahseers and their habitat, promote Wildlife Value Orientations (WVOs) of mutualism and respect towards biodiversity (Fulton et al. 1996; Gomez et al. 2022) leading to greater pro-conservation activities. Furthermore along with adding evidence-based scientific inputs to the plans and protecting interests of each stakeholder, such a system will ensure shared accountability for conservation problems and failures as well (Pimid et al. 2022). Interestingly, examples of multi-stakeholder collaborations for mahseer conservation with the potential to replicate in other parts of the country were also recorded from selected regions of Karnataka. The joint initiatives between the recreational anglers and fisheries department in Kodagu (25), partnerships between anglers and forest department in Mandya and Chamarajanagara (24) and the ongoing collaborations between researchers and fisheries department (12) are noteworthy.

**Fig. 6.**
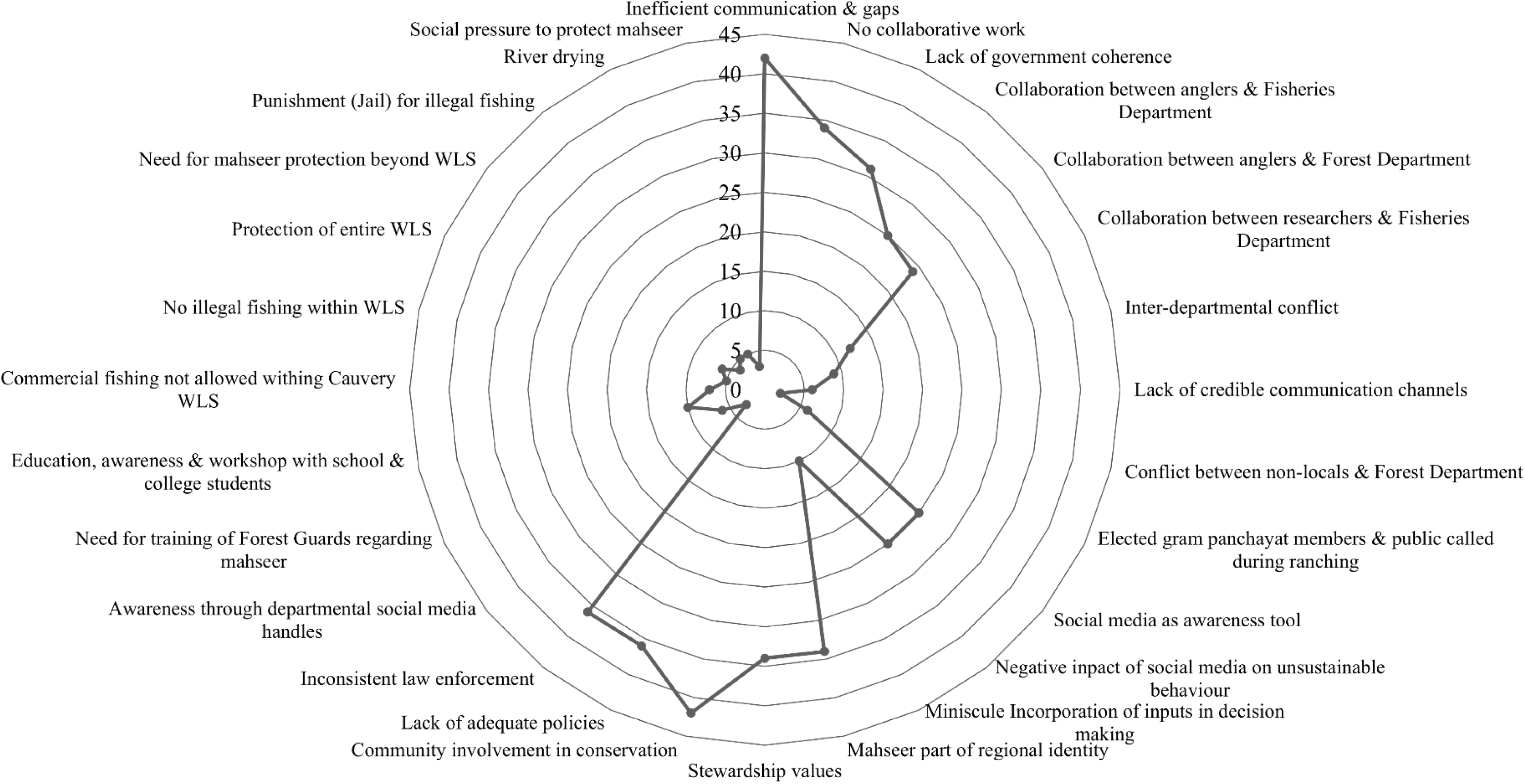
Radar chart depicting the frequency of codes used for generating the theme ‘communication and collaboration’ from the reflexive thematic analysis conducted on the interviews and FGDs from Karnataka.

The collaboration between different stakeholders at micro level can be initiated by involving villagers and members of the local governing bodies such as *gram panchayat* (6) during fingerling ranching, post release monitoring, angling activities and engaging former poachers as river guards or *ghillies* etc. These actors demonstrated strong intrinsic and relational values, often driven by region-based pride, regional identity and cultural connection to the mahseer and the river (34) alongside their pronounced stewardship values (34). Such a relational view of the species offers a powerful foundation for conservation, fostering respectful behaviour towards wildlife (Brackhane et al. 2019; Jolly and Stronza 2025).

> Quote 4: *“The local community members who have been fishing for generations have immense local knowledge about how to sustainable coexist with wild animals. They respect and value the animals, the water, the river. Everyone should learn from them. The decision makers should incorporate their views and knowledge, involve them in decision making discussions for effective management”.*

Insights from TPB affirm that when attitudes and social norms are aligned, and behavioural control is adequately supported, participation and conservation behaviour can be sustained (Ullah et al. 2021; Karimi and Ataei 2022). Hence, people are more likely to participate in conservation and management activities if their knowledge and inputs are sought, incorporated and built upon highlighting their importance to conservationists and officials (Pretty and Smith 2004; Young et al. 2013; Setchell et al. 2017). However, such participation remains extremely under-recognised in the context of mahseer conservation. Incorporating local community members and their knowledge in decision-making (42), community involvement acknowledging cultural contexts (e.g., religious-significance based site selection for temple-based *matsyadhamas*), and offering both monetary and symbolic recognition (through certificates, news media acknowledgments, etc.) can enhance long-term engagement and partnership between different actors involved in mahseer conservation and the people inhabiting such areas.

The theme ‘values and moral responsibility’ along with its sub-themes, stewardship (34; relational; Fig. 7), socio-cultural association and meaning (31; relational) and mahseer as part of nature (42; intrinsic), is associated with the all 3 components of TPB, social values and the second and third phases of CPF. The recreational anglers, especially the multi-generational ones, exhibited strong relational values, viewing mahseer as a natural heritage (25) and as part of their identity (25) with their lives being intertwined with them (25; all relational).

> Quote 5: *“We feel that it is our duty to protect the mahseers, maintain a healthy aquatic ecosystem, monitor any illegal activities, make people aware about the benefits of sustainable angling and make sure that the species survive for our next generation to angle, just like we learnt from our previous generation. Such activities give us a sense of peace, happiness and calm.”*

**Fig. 7.**
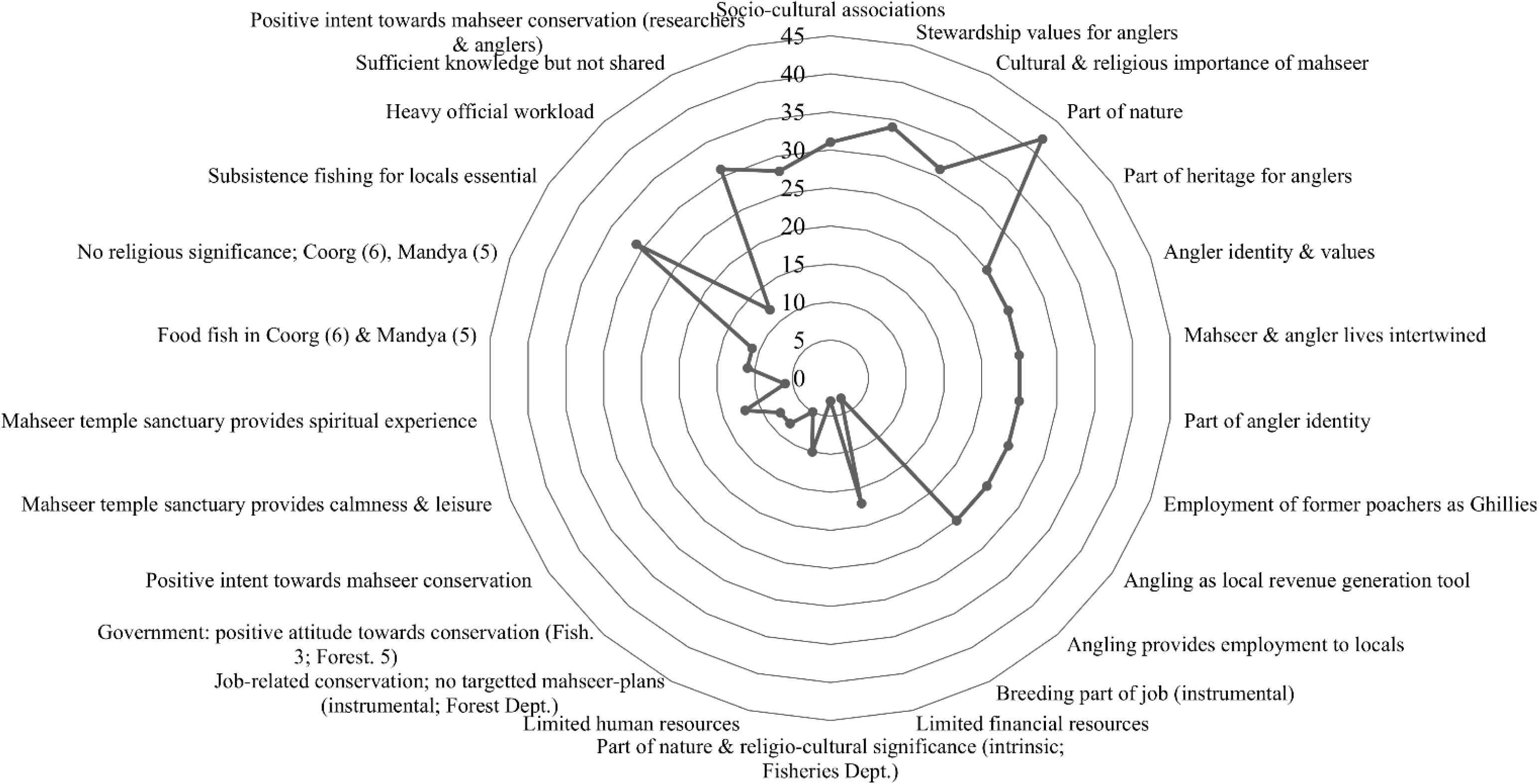
Radar chart depicting the frequency of codes used for generating the theme ‘values and moral responsibility’ from the reflexive thematic analysis conducted on the interviews and FGDs from Karnataka.

Amongst the anglers, the shared identity (25) of being the caretaker and protector of the river and mahseers, hence the peer pressure from the community members may potentially be fostering a culture of voluntary stewardship and motivating them to participate in patrols, catch-and-release fishing, and reporting illegal activities, employing former poachers and locals as river watchers or *ghillies* (25) and involving local community members in such activities thereby contributing to the local revenue generation (25). However, the staff from the fisheries department expressed both intrinsic and instrumental values towards mahseer. They considered the fishes as an integral part of nature and acknowledged their religio-cultural significance (3; intrinsic), while simultaneously reflecting instrumental values linked to their professional responsibility through active participation in conservation breeding initiatives (3). The forest department staff similarly demonstrated job-related instrumental values associated with broader wildlife protection (5), although they did not possess any targeted mahseer-specific conservation approach. Such values may have resulted in their intent towards conservation being slightly less positive (3; fisheries, 5; forest).

> Quote 6: *“We are very understaffed, we also have a lot of official work, paperwork to complete and other government duties beyond mahseer conservation, so we cannot completely invest our time in this. It would be good both for us and the mahseers, if there were a dedicated mahseer conservation unit with more staff and funds”.*

This statement from a fisheries department staff provides insights on how both the ABC and PBC can limit conservation efforts. Although the government staff (both fisheries and forest department) demonstrated a positive attitude towards conservation (8), shortages of human (10) and financial resources (17) may reduce their government staff’s perceived and actual ability to act. Additional barriers such as heavy official workload (12) and unshared available knowledge (31) can further serve as a barrier to conservation efforts. Such misalignment between intent and capacity highlights the need for incorporating TPB into CPF implementation and monitoring to understand if the prevalent enabling conditions are addressed alongside attitudes and social norms. A positive attitude, with supporting social pressures along with enough perceived and actual behavioural control has been shown to lead to a positive intention of performing behaviour (Ullah et al. 2021; Karimi and Ataei 2022).

Mahseers are attributed with a high level of socio-cultural values and divinity (relational values) particularly, at the Ramanathapura temple-based fish sanctuary area in Hassan district. Local people irrespective of gender, age and occupation come and pray in the temple first; then they move to the river bank and pray to the water and the large aggregation of mahseers present there. They offer them *prasada* (offerings consisting of flattened and puffed rice, peanuts, and flowers) as an act of devotion and spend time by sitting by the river banks, reflecting the relational values that the local devotees attribute to these fishes.

> Quote 7: *“These fishes are God for us. We pray to them and make wishes. No one is allowed to catch and eat them here”, one of the devotees we met said. The people visiting this sanctuary claim to achieve calmness (12) and spiritual experience (6; relational) by watching, praying and feeding the fish, stressing on the non-tangible cultural ecosystem services (CES) provided by them.*

The mahseers also contribute also to the local economy. Quote 8: “*Our livelihood depends on the devotees, who buy puffed rice, flowers, and other offerings from us to worship the mahseers.*” However this divinity enjoyed by the mahseer is not uniform in the entirety of Karnataka. In Kodagu and Mandya districts, these fishes are perceived primarily as game and food fish (instrumental) with no religious significance attributed to them by the local community (6, 5 respectively; Fig. 7). Such discrepancy stresses the need for considering local cultural traditions while designing programmes for mahseer conservation.

### Assam

This state also actively implemented multiple programs to conserve their mahseer populations - captive breeding in hatcheries (52; Fig. 8), community-involved awareness programmes (41), ‘Mahseer Watching’ initiatives (10), imprisonment for poaching within protected areas (30). However, CPF (situation assessment phase) disclosed defunct hatcheries (45) and ineffectiveness of ‘Mahseer Watching’ (6). ‘Consensus between stakeholders’ reappeared as a major theme in Assam as well. Noticeably, all the stakeholders we met had a positive attitude towards mahseer conservation (114; TPB; Fig. 8) in this state during our study. The concurrence on the decline of mahseer in Assam (66), and siltation and river-bed mining (68), destructive fishing methods (59), and commercialisation of traditional community fishing practices (67) as the major threats for these fishes, and the urgent need for implementing conservation measures (66) were also noticed. Many shared concerns over the decline in the native fish species (53) and increase in IAFs (53) caught during the community fishing events in the recent past. Furthermore most of the researchers and anglers (17) we contacted, were of the opinion that IAF such as common carp, grass carp, and silver carp are quickly growing as the competitors for the natural populations of mahseers in this state. However, competing stakeholder interests on economic development versus conservation, and the jurisdiction issues existing between different governmental organisations involved in conservation of mahseer may hinder the smooth implementation of such programmes. For instance, the majority of respondents from the fisheries department were primarily interested in livelihood generation and aquaculture-related activities (24), while anglers, researchers and local community members were vocal about conservation needs. The words of an aqua-tourism centre manager vouch for promoting stakeholders consensus as soon as possible.

> Quote 1: *“We have told the officials multiple times that the beel can be used rationally, some for aquaculture related breeding of IMCs and other economically important fishes, while the others can be used for mahseer and other endemic fish conservation breeding.”*

**Fig. 8.**
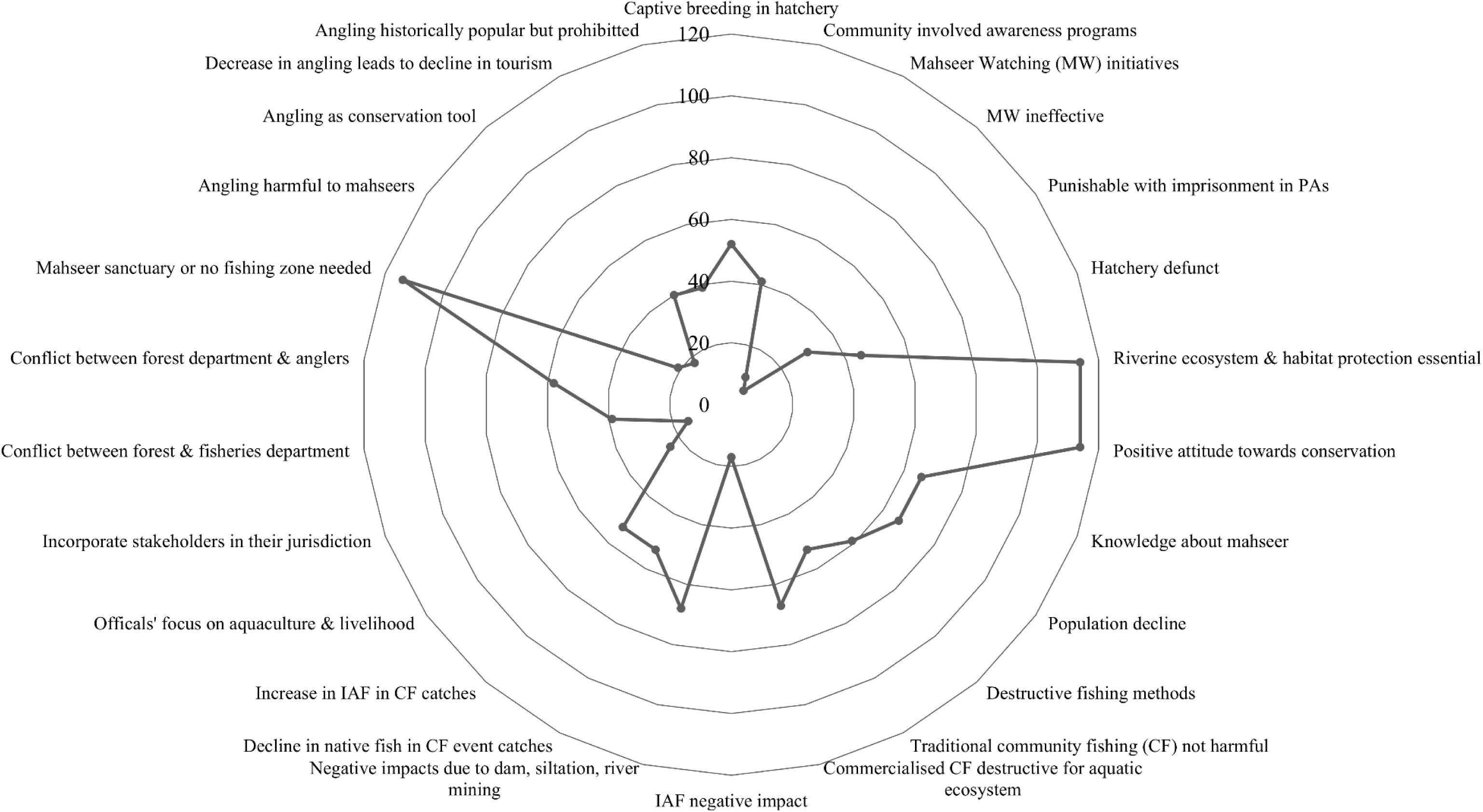
Radar chart depicting the frequency of codes used for generating the theme ‘consensus between stakeholders’ from the reflexive thematic analysis conducted on the interviews and FGDs from Assam.

Similarly, forest department staff expressed their displeasure on the exclusion from mahseer breeding and restocking activities undertaken by ICAR-DCFR and anglers within areas under their jurisdiction (15). In contrast, anglers believed that the activity of production of mahseer juveniles and releasing them into natural waterbodies does not come under the responsibilities of the forest department, and hence it is not mandatory to involve their staff in such activities.

The forest department having tension between fisheries (39) and the former and anglers (58) were also not rare.

> Quote 2: *“Eco-camp once supported mahseer breeding with anglers and scientists working together. The forest department was however, not involved in such programmes despite the Eco-camp being located within the buffer zone of the National Park and right next to the DFO office. This exclusion may have angered the forest staff. As much as we support the forest department’s decision to ban angling in protected areas, the closure of the camp led to decline in foreign tourists and anglers, which affected our income and many villagers who were employed there, lost their jobs.”*

This statement made by a local indigenous community member reflects the second theme generated, ‘communication and collaboration’. In Assam, the decision-making and governance system guiding mahseer conservation was largely top-down, with lesser opportunity for meaningful incorporation of local, indigenous values, and scientific insights. Political influence, bureaucratic and legal impediments, and fragmented inter-departmental responsibilities were found to play critical roles in reducing the sustainability of the efforts for maintaining collaboration amongst the stakeholders. Communication gaps (67; Fig. 9) and very little interaction (51) existing between the forest and fisheries departments often resulted in the desynchronisation of the mahseer conservation activities (53) conducted by these two pillar institutions. The disconnect, bureaucratic pressures and legal challenges, the barriers of collaboration and cooperative decision-making (43), left the hatchery and eco-camp present in the Nameri National Park jointly run by ICAR-DCFR, anglers (15) and state fisheries department (16) defunct depriving the local community members of both revenue and livelihood opportunities from eco-tourism. According to major stakeholders (organisations involved in the management of the lakes) of Mahseer Watching initiative by the ICAR-DCFR in the *pukhuri*/lakes across cities such as Guwahtai also failed because of poor planning (6), and limited human resources (6) resulted from the lack of consultation with the people working on the ground.

> Quote 3: *“Mahseer watching was not well thought-out. Since fish were invisible, the public were not interested to visit. The officials did not think of that nor did they ask us. We have only a couple of staff managing the entire pukhuri and the park, no one thought of who would take care of the fish, feed them” - said one of the local pukhuri management staff members.*

**Fig. 9.**
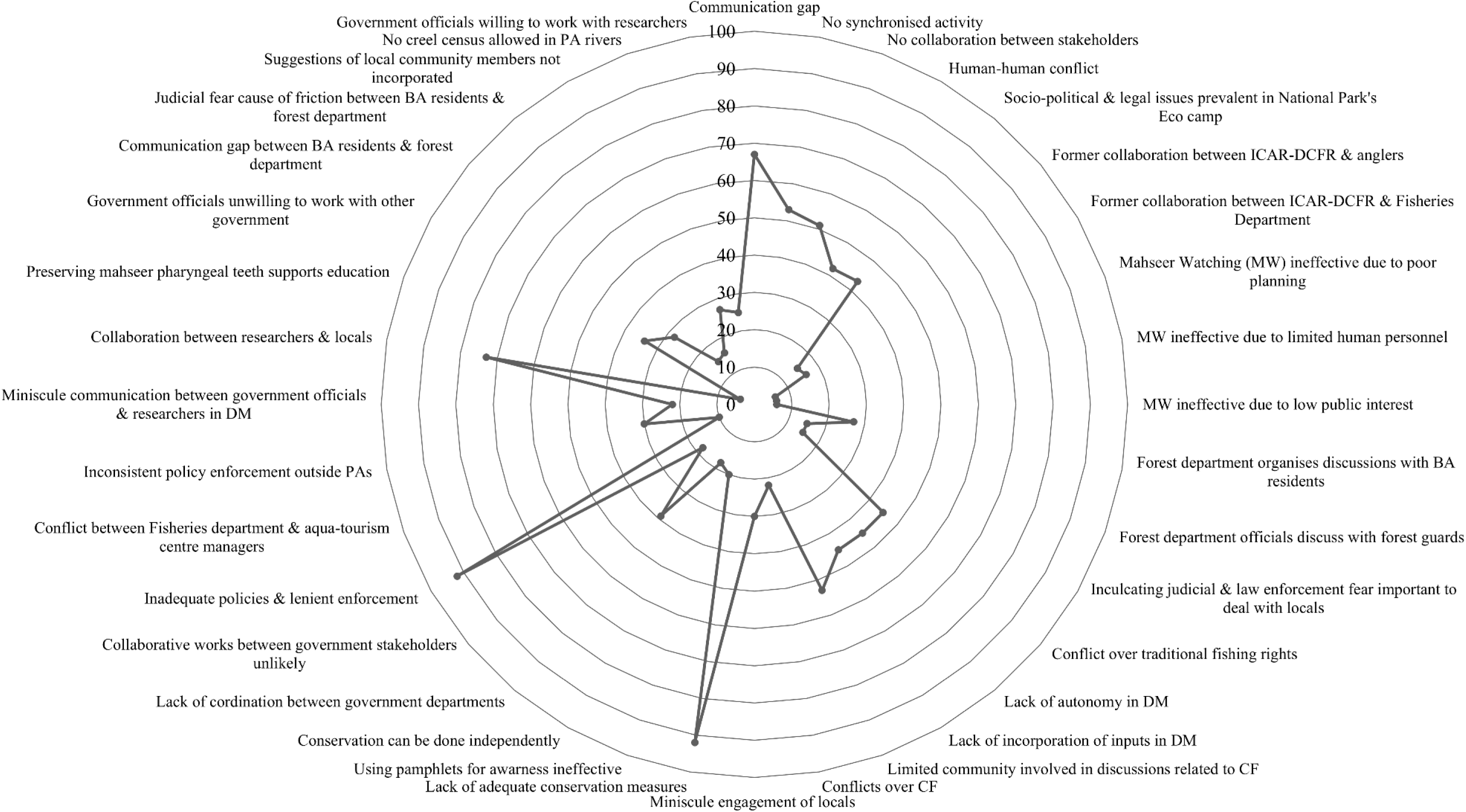
Radar chart depicting the frequency of codes used for generating the theme ‘communication and collaboration’ from the reflexive thematic analysis conducted on the interviews and FGDs from Assam.

Another group of people who suffered from the unavailability of a trustworthy mechanism to answer their queries and guide their fishing activities were the buffer zone communities. Although forest officials acknowledged periodic discussions with these stakeholders (27), the laypersons unaware of the legal dimension of the fishing in the buffer zone were much concerned about the punishment (imprisonment) for subsistence fishing in their localities (15). Furthermore, interactions between forest officials and buffer zone communities seldom transformed into substantive decisions or had the potential to influence the policies (45). Similarly, diminished interactions and joint activities between government agencies and local community members (53) weakened public trust and introduced frictions (22) in the community fishing events. Hence, acknowledging lived experiences and knowledge of the locals by the authorities, and involving them in the conservation activities along with installing an effective system for bidirectional communication can reduce the resistance from the communities inhabiting landscapes important for managing mahseer populations.

The stakeholders expressed a mix of intrinsic, relational and instrumental values towards mahseers (theme 3: ‘values and moral responsibility’). These fishes were considered as an integral part of the nature (69; intrinsic; Fig. 10), food source that has been historically fished for generations (88; instrumental), especially *N. hexagonolepis* (66), and many individuals we contacted, felt a moral responsibility to protect them (83; relational). The long-standing socio-cultural connections with mahseer (relational values) echoing the strong pro-conservation social and subjective norms (TPB) prevalent within the community (109) was clearly visible in the words of an indigenous community member:

> Quote 4: *“Mahseers and all other animals are a part of nature; they have their own right to live. Our lives are intertwined; we have traditionally fished them for generations as they are a food fish for us. It is our duty to safeguard these fishes.”*

**Fig. 10.**
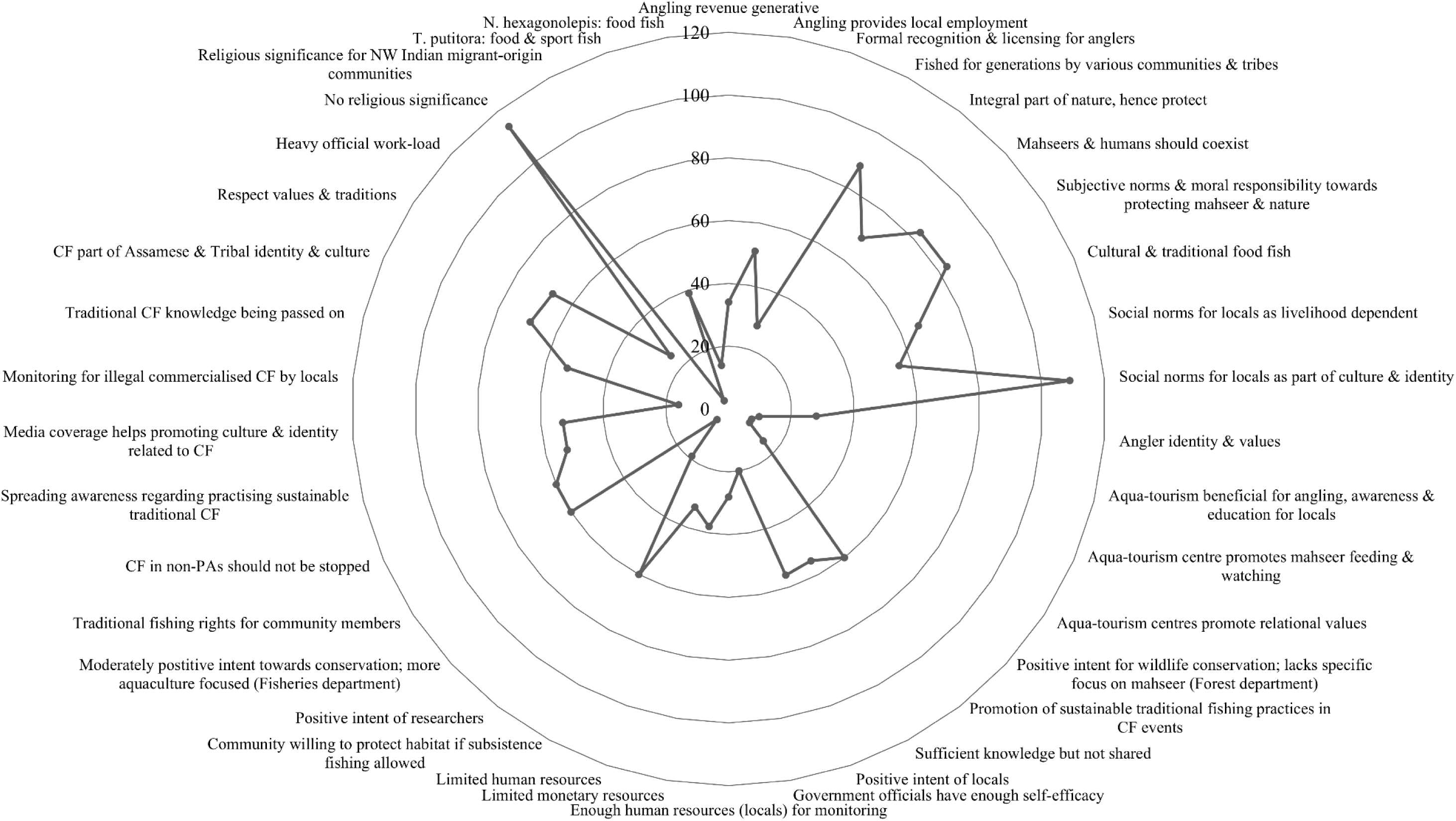
Radar chart depicting the frequency of codes used for generating the theme ‘values and moral responsibility’ from the reflexive thematic analysis conducted on the interviews and FGDs from Assam.

Anglers community were also vocal about their activities for promoting the practice of C&R angling, fostering voluntary stewardship and monitoring illegal activities, and the role of caretakers of the rivers and mahseers (28). Many anglers suggested engaging local community members in angling-related eco-tourism activities to generate revenue (34) and employment (51) at the local level. Aqua-tourism managers also advocated promotion of ethical C&R angling, raising public awareness and education (10), and in their opinion offering opportunities for the visitors to interact with, feed, and spend time with mahseers, and introducing the ecosystem services received from them (8) may catalyse the inculcation of relational values (8) towards these fishes among the general public.

The government officials from different departments acknowledged the importance of biodiversity conservation, but a domination of instrumental values over the relational aspects was noticed in their responses. Furthermore, in many contexts the species-specific conservation intent for mahseer (e.g. forest staff - 15) was also not conspicuous. Although, a high level of behavioural control is attributed to these stakeholders by others, the dearth of monetary (38) and human resources (33), human-human conflicts (42), heavy official work-load (25), reluctance for inter-departmental collaboration (34), poor knowledge sharing mechanism (55), and restricted autonomy in decision making (45; ABC) impacted the translation of their belief in self-efficacy (20) into the action in the form of better governance, monitoring and evaluation (CPF). However, the local community members with traditional fishing knowledge displayed strong relational values and willingness to protect mahseer and their habitats (60). Motivated by the stewardship norms rooted in their cultural identity (109) and livelihood dependency (56) on the immediate environment they claimed to have sufficiency in the human resource (PBC; 28) required for monitoring mahseer habitats if traditional fishing rights are recognised (60).

> Quote 5: *“The government should allow subsistence fishing for the community members; they have traditionally done it for years without harming any species. If you allow them to do this, they will themselves monitor the river stretches preventing illegal activities and poaching.” These words of an angling association member working closely with the local indigenous community, points towards the need for utilising their familiarity with the focal environment and experience for effectively managing the mahseer population. However, enforcement of rigid policies and lack of recognition of traditional fishing rights of the buffer zones communities and ban on subsistence fishing, which limited their livelihoods and ignited resistance and friction with the authorities (45). Anglers, another stakeholder group with strong relational values, moral responsibility, and pro-conservation intent, and involved in monitoring mahseer habitats and promoting eco-tourism historically in this state also complained about restrictions and absence of a mechanism issuing formal licensing (28).*

Involving representatives of the local population in the organization and conduction of community fishing events (53) could enhance the quality of governance and monitoring, and can also generate opportunities for eliminating an important misconception kept by many - the traditional fishing gears are not ecologically harmful (53). If safeguarded from commercialisation, community fishing festivals such as *Junbeel Mela* could serve as a powerful platform for promoting moral responsibility, cultural identity, social values and conservation-positive subjective norms of coexistence (83), alongside Wildlife Value Orientations (WVOs) of mutualism and coexistence (Fulton et al. 1996; Gomez et al. 2022). The following statement of a community member reflects this statement.

> Quote 6: *“Community fishing is deeply etched in me as my culture, my identity, my heritage. We are very proud that we are able to promote cultural community fishing by inviting delegations from different states of India each year. Sadly, I have seen people illegally catching fishes with non-traditional gear, that is wrong and we do not support it. We have always practised non-destructive fishing for generations. It is also sad to see people making bandhs/bunds in the river channels feeding to our beel as they also affect the fish populations.”*

However, local indigenous community members felt that commercialisation and destructive fishing practices (67), driven by profit motives and inadequate regulation, have eroded the relational values despite the inter-generational transmission (53) of the knowledge, and their exclusion from the vital roles of community fishing (53) restricted them from working for eliminating the human-human conflicts (22; Fig. 10) associated with such events. These results demonstrate how a top-down approach in governance without acknowledging the values, feelings and knowledge of the local communities and stakeholders can reinforce behaviours counterproductive to conservation objectives (Dhee et al. 2019; Jolly and Stronza 2025; Riechers et al. 2025).

### Uttarakhand

Similar to Karnataka and Assam, several mahseer conservation initiatives such as enhancing public awareness through street plays, dance, skits, radio programmes, local newspapers (8; Fig. 11), community involved communication programs (16), and imprisonment for illegal fishing within protected areas (8), were observed also in Uttarakhand (situation assessment – CPF). However, this state gave a greater emphasis for mahseer ranching, a hatchery dedicated for these fishes run by the state Fisheries Department in association with ICAR-DCFR at Bhimtal, Nainital, undertaking captive breeding in semi-natural conditions to increase the survival of juveniles post-release (8) and the relocation of fingerlings trapped in river stretches that dry up during the summer to safer upstream sites (3) by the fisheries department are a few examples.

> Quote 1: *“We are very much invested in the survival of the fingerlings. During the summer months, the middle and lower stretches of many rivers dry up and the small fishes from our earlier release get trapped in the puddles of water. We pick them up from these puddles and release them upstream where there is sufficient water” – said a fisheries department staff, reflecting the value given to the fingerlings by the authorities in this state.*

**Fig. 11.**
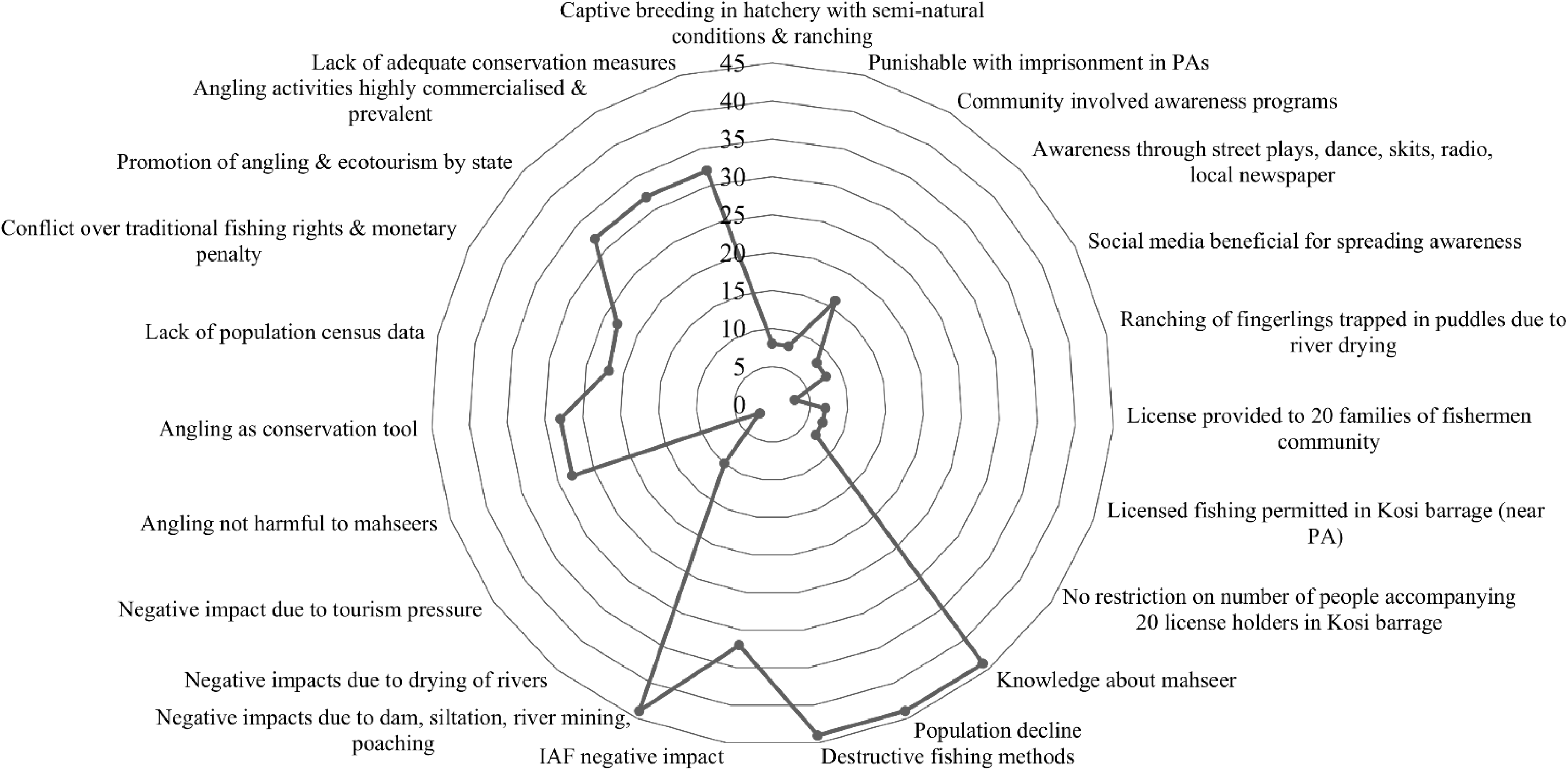
Radar chart depicting the frequency of codes used for generating the theme ‘consensus between stakeholders’ from the reflexive thematic analysis conducted on the interviews from Uttarakhand.

However, the situation assessment (CPF) revealed many inefficacies in the conservation programmes induced by the fragmented regulation and monitoring systems, uncoordinated activities by the stakeholders, inadequate inclusion of diverse groups with interest in the ecosystem management, loopholes existing in the mahseer fishing licensing and an indolent state-wide mahseer-specific conservation interventions. For instance, although a fixed number of individual fishing licences are officially issued annually for the Kosi Barrage in Ramnagar, Nainital (7) for the period from November to June, there is no restriction on the number of individuals (7) who may accompany the licensee to the field. This ambiguity is often misused by the parties with vested interest resulting in excessive fishing pressure, as revealed by a forest official.

> Quote 2: *“Recognising the livelihood needs and for the welfare of the fishing community, we have allowed fishing in the Kosi barrage area, which lies along the border of the Jim Corbett National Park buffer zone. We issue 20 licences to fishermen community members for a fee to fish, sell their catch and eat. But the problem is, there is no restriction on how many people can accompany a license holder, so the pressure on the fishes has increased. There is no proper mechanism to monitor what is happening there as well.”*

Almost all stakeholder groups we met agreed (‘consensus between stakeholders’ - Theme 1) that mahseer populations are critically declining in the wild (44; Fig. 11), and they demanded urgent interventions for the conservation (44). Along with throwing light on the reasons behind such a reduction in the population size, sub-themes generated - dam construction, siltation, riverbed mining, poaching (44), absence of population census efforts (22) and ecological disturbances caused by invasive alien fishes (IAF) - also reiterated the need for immediate collective actions. Although the presence of IAF common carp (32) was perceived as a threat for local fishes, none of the stakeholders acknowledged the impact of another invasive fish tilapia spreading in the waterbodies of Uttarakhand (Trivedi et al. 2021). Both officials from the forest department and NGOs agreed on drying up of the rivers (10) due to the indiscriminate extraction of water by hotels (pressure from the tourism) and houses impacting mahseer populations. Interestingly, many focal groups viewed recreational angling, a key eco-tourism component, not as a threat to the mahseers but a potential mechanism for ensuring their protection (28) pointing towards the possibility of merging conservation goals with the livelihood of the local people to gain public acceptance for the conservation interventions.

Another context in which lack of consensus between the two important stakeholders, fisheries department and local people led to the conflict was the fine for illegal fishing (23). According to a staff of the fisheries department:

> Quote 3: *“The fine for illegal fishing initially was Rs. 2,000 with jail time, but locals found it too high, which led to anger and conflicts. So, we decreased it to Rs. 500 with no jail. However, the instances of illegal fishing have increased over the years, so I think we will enforce stricter measures again. While we support traditional subsistence fishing through daily fishing licenses for locals, we strictly discourage large-scale illegal fishing that harms the ecosystem. As a result of such mass scale killing, we are not getting enough brood stock for hatchery breeding”*

These words should be taken seriously by the authorities and a consensus on the punishments for illegal fishing should be reached as soon as possible after conducting proper stakeholder consultation to avoid further escalations. Establishment of a platform under the direction of the institutions of local governance for promoting interaction between the stakeholders will also help in reducing the friction between government departments (forest and fisheries department, 16), forest department and non-license-holding locals (8), fisheries department and locals (7), and locals leaders and NGOs (7) reported from the study area.

The attitude towards mahseer conservation was positive (44; Fig. 12) in this state also, but limited knowledge sharing (32) and collaboration (32), lack of coordinated efforts between government departments (32), and disjointed and limited communication (32) (Theme 2 - ‘communication and collaboration’) restricted translation of it into actions on the ground. For instance, though an informal collaboration between ICAR-DCFR and the fisheries department existed (7) no efforts to formalise it, which may catalyse the joint activities, was active. Furthermore, failure to incorporate vital feedback from key actors into decision-making or planning processes (10), inconsideration of the inputs from the researchers (4) and advises of the NGOs (10) point towards a top-down approach prevalence in the mahseer conservation planning and interventions in the study area. The efforts in the past few years to involve local communities or local Self-Help Groups (SHGs) by leasing selected river stretches of 5 to 8 km, under a fee-based beat system (11) have not yielded the expected positive outcomes. Limited communication between the government departments and local beneficiaries, the absence of a clear statement of the purpose and expectations of the leasing and failure to implement a mechanism to monitor the activities of leases (8) declined public interest (8) in this initiative. This situation rendered local participation largely income-driven rather than conservation-oriented, stressing the need for maintaining regular discussions between authorities and non-government organisations, as denoted by a member of a local nonprofit organisation working for the mahseer conservation.

> Quote 4: *“After proper communication only the fisheries department should lease the river beats to NGOs who are accountable and interested in conservation and protecting the rivers along with sustainable fishing. There was no long-term collaborative effort from either the officials or the locals, hence people started losing interest. Many local groups who were leased the river stretch did not undertake any monitoring or safeguarding of the rivers which indeed led to more unsustainable fishing.”*

**Fig. 12.**
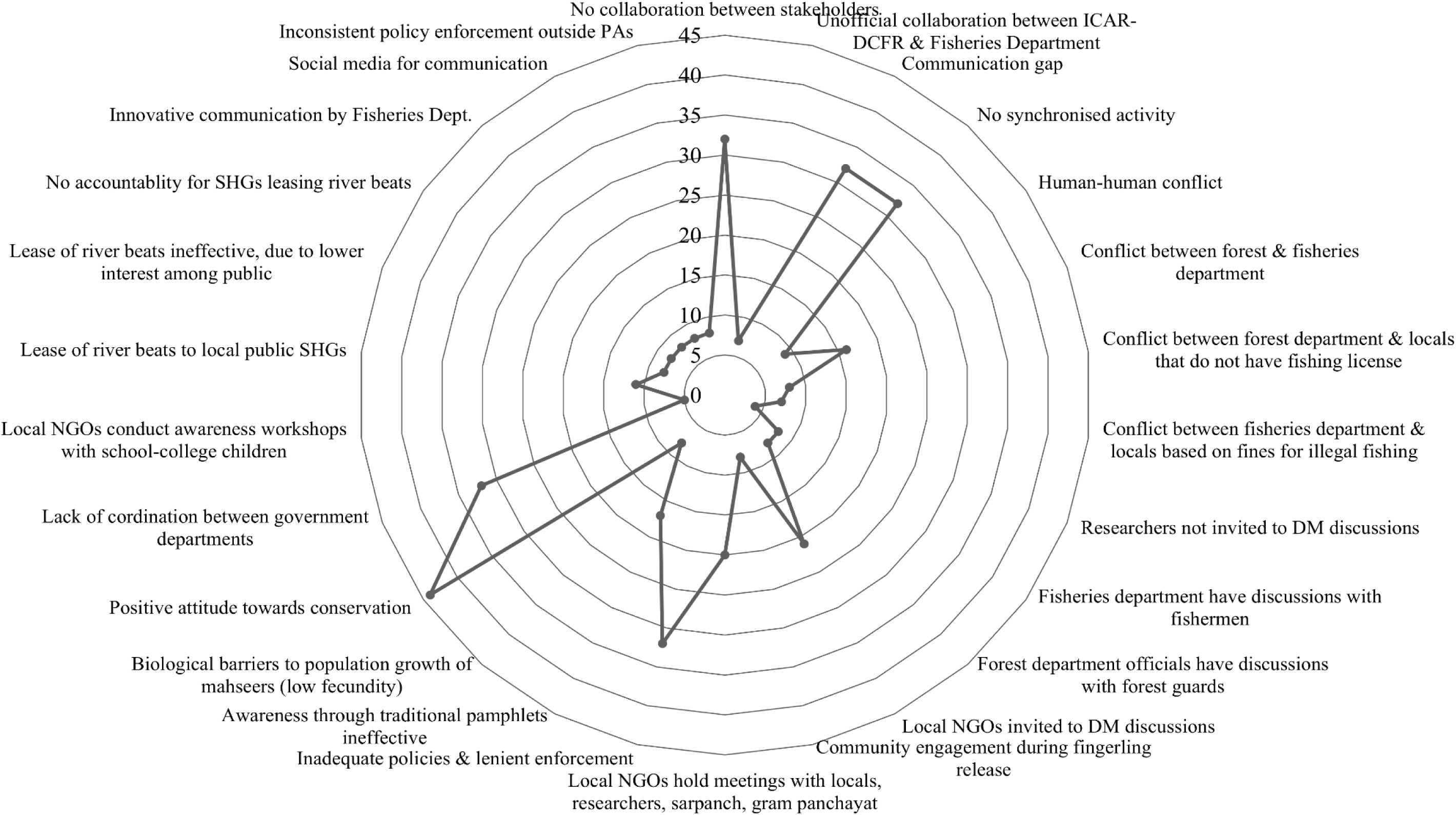
Radar chart depicting the frequency of codes used for generating the theme ‘communication and collaboration’ from the reflexive thematic analysis conducted on the interviews from Uttarakhand.

The forest department staff often consulted forest guards (8) and the fisheries department engaged with fishermen (8), but outcomes of these interactions were seldom considered for co-management and collaborative planning. However, the fisheries department staff conducted many innovative public awareness and communication programmes such as skits, *nukkad natak* (street plays), etc. They used local radio channels and newspapers (8) and social media (8) as the medium for information dissemination.

> Quote 5: *“We have been trying to spread awareness about mahseer decline and conservation in innovative ways. The traditional pamphlets and newsletters do not work; no one reads them. We are organising nukkad natak, planning to display slogans and hoardings, play songs on radio and use social media. Making things go viral is the best way to spread awareness. We plan to take this further by making humorous graffiti and street comedy stand-ups in local languages. These have more impact on the locals.”* - said a staff member of the fisheries department. Local NGOs were also active in conservation education and community outreach. They conducted workshops and community meetings keeping school children and college students (5), village sarpanch, gram panchayat members, and locals (20) in the focus.

The theme ‘values and moral responsibility’ emerged also in the state of Uttarakhand. Local communities and NGOs viewed mahseer as an integral part of nature (17; intrinsic; Fig. 13), symbol of culture and identity (17; relational), and a source of food and livelihood (17; instrumental). Their belief, protection of this fish is their moral responsibility (17; relational), was echoed as strong pro-conservation social and subjective norms (TPB) prevalent in the study area (17). Here, youth displayed particularly strong moral responsibility and supported conservation-positive norms (10). However, members of the local community expected permission for subsistence fishing (12) and discount in the monetary penalties for illegal fishing in return for the mahseer protection (9) since fishing was a major source of livelihood for many (12). Perceived behavioural control (PBC) for monitoring of mahseer habitat (human resource; 17) was high for both local communities and the NGOs but the latter complained about the political pressures (5) they face.

**Fig. 13.**
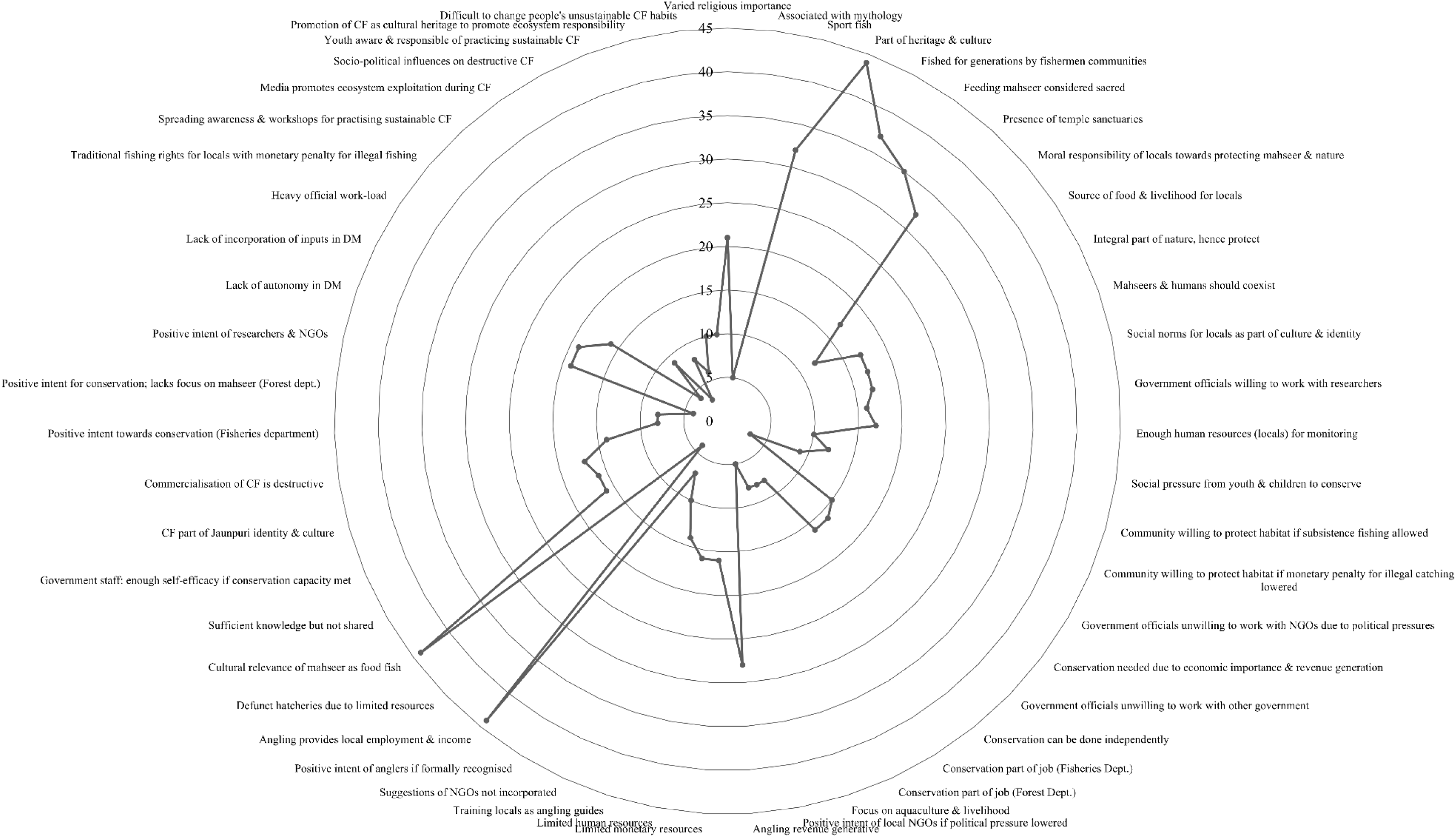
Radar chart depicting the frequency of codes used for generating the theme ‘values and moral responsibility’ from the reflexive thematic analysis conducted on the interviews from Uttarakhand.

Anglers, another important stakeholder, expressed instrumental and relational values and positive conservation intent but felt that authorities do not recognize and value their activities (7).

> Quote 6: *“Angling is hugely beneficial for the entire village. We share a major proportion of revenue earned from the visiting anglers with locals, train youths as angling guides, and buy all our vegetables from the local produce. It is sad that we do not have any recognition as anglers or guides. In Bhutan, anglers receive formal licenses or identity cards; we also want the same respect and legitimacy. We also request the Fisheries Department to set up a local office near angling hotspots, to partner with us and help us monitor and prevent illegal fishing.”* - said a member of the angling association.

Along with anglers, multiple other stakeholders were also of the opinion that regulated and formalised, angling can generate revenue (28), create local employment opportunities (44), engage and train youth as angling guides (14), and support conservation through license fees and creel census data. As a result, the anglers expected the governance and decision making mechanism (CPF) would implement a formal licensing system in the near future and promote partnership between the fisheries department and the anglers to monitor the fishing and angling activities. They also expected that such a collaboration along with reducing the conflicts, governance gaps, could promote eco-tourism benefiting local communities and generate supplementary conservation funds (Phil 2022) to resolve bottlenecks induced by the limited government grants (Echols et al. 2019) available for mahseer conservation.

In Uttarakhand, particularly in regions such as Bhimtal in the Nainital district, the mahseer hold significant religious (21) and mythological (5) importance (relational value), forming an integral part of the local folklore. In Uttarakhand, particularly in regions such as Bhimtal in the Nainital district, the mahseer holds significant religious (21) and mythological (5) importance (relational value), and this fish is an integral part of the local folklores. According to one such legend, the *Nal-Damyanti Tal,* a prominent lake located in Bhimtal with a mahseer population is believed to have emerged at the very site where the palace of mythical King *Nala* and his consort *Damayanti* once stood. Lord *Indra*, the King of Gods in Hindu mythology, collapsed the palace into a lake while cursing the royal couple. Later when *Indra* reversed the curse the lake came to be filled with golden mahseer, infused with *amrit,* the elixir of life. Myths hold that when the starving King *Nala* once tried to roast a mahseer, it leapt back into the lake, though with a mark of the fire on its body, proclaiming the boon of ‘*amrit*’ and immortality. Revered as eternal beings and guardians of the lake’s sacred essence, the mahseer of *Nal-Damyanti Tal* are venerated. Fishing in this lake is strictly prohibited and for locals harming mahseer means hurting the Gods themselves (Kattyayani 2016; Vishwakarma *personal communication*). Mahseers are revered in several other non-formal temple fish sanctuaries (32) also in this state, such as in Baijnath temple, Garjiya Devi temple located on the shores Gomti River and Kosi River respectively, where feeding the fish is considered to be sacred (35). These temple sanctuaries offer protection to mahseers though the religious significance varies across communities.

Far more than catching fish, community fishing events such as the *Maund (Maun) Matsya Mela* held in the Jaunpur region of Tehri Garhwal keeps a deep connection with local culture, Jaunpuri identity, and heritage (17). Participating in this century-old annual festival, initiated by *Raja* Sudarshan Shah of Tehri in the mid 19th Century is regarded as an ancestral right (Sharma et al. 2016) by the Jaunpuri community. Preparations for the ‘*mela*’ begin one to two months in advance, and nearly ten to fifteen thousand villagers from Jaunpur (Tehri Garhwal) and Jaunpur-Bhabar (Dehradun) areas, attend this fishing festival underscoring its religious, cultural and historical significance (Sundriyal and Kumar 2019; Rajpoot *personal communication*) and its role in strengthening social bond and cultural integrity among the participants (Singh et al. 2016; Sundriyal and Kumar 2019). However, a range of contemporary issues such as limited institutional recognition and reinforcement, shifting local priorities, and increasing migration have shaped this event in recent years (Habib 2025). As a result, what was once a community-bonding ritual and celebration has gradually become commercialised and destructive (14) in the recent past, triggering human-human conflicts and eroding relational values, as reflected in the words of a local NGO member:

> Quote 7: *“The traditional Maund Mela has now become completely commercialised, the essence of the community belongingness, culture and tradition is not there anymore. Nowadays, people use chemicals, poisons and nets in the river, play loud music, have a party and throw garbage in the river. Sadly, due to very high political influences and pressures, no one says anything. Many NGOs are raising awareness but it will take time. Positively, some of the youth have shown enthusiasm and they refrain from such activities.”*

Stakeholders also complained that socio-political factors, including political influences and power-dynamics led to politicians supporting such destructive fishing (8) during the *Mela*, and the media reinforcing this negative trend by projecting destructive events rather than highlighting ecosystem degradation and its implications (3). On a positive note, the NGOs and researchers (also supported positive conservation behaviour; 4) have been working to restore relational values among locals by conducting awareness campaigns and workshops on sustainable community fishing (9). Even though interest in conservation and moral responsibility are gradually increasing among the local youth (6), changing practices of the community may take a long time (10). Therefore, promoting regulated community fishing using traditional gears and recognising these events as socio-ecological heritage and a catalyst for eco-tourism, engaging local youth, Jaunpuri community members, and NGOs in overseeing it, leveraging youth-elder relationships and encouraging news media to highlight sustainable, culturally grounded practices which form an integral part of Jaunpuri identity and culture (17) is a need of the hour to foster ecosystem responsibility (10).

Staff from different government departments varied in the values attributed to mahseer. Fisheries department staff expressed concern over mahseer decline and actively participated in conservation programmes. Their job-related instrumental values (8) were clearly evident in their focus on linking aquaculture with livelihood (8) for the locals. Many acknowledged cultural and heritage significance (relational; 8) that mahseers have in their area and a positive intent (8) to protect them. Officials from the forest department were also committed to the protection of biodiversity (job-linked instrumental values; 8) and recognised this fishes’ cultural importance and the connection with local identity (8; relational), but were not vocal about mahseer-specific conservation plans like the members of fisheries department. Nonetheless, staff from both departments were more open to work with researchers (16). Although both departments reported self-efficacy (PBC 16) for mahseer conservation they also admitted the constraints faced during ground action in the form limited finance (16), human resources (16), heavy workloads (16), restricted decision-making autonomy, (16) and poor knowledge sharing (16; Fig. 13). This misalignment between PBC and ABC could hinder effective governance and monitoring and hence the conservation outcomes.

## 4. Discussion

The present study revealed several commonalities in mahseer conservation across the three geographically and socio-culturally distinct regions of Southern, North-Eastern, and Northern India, while highlighting local initiatives holding potential for replication to safeguard aquatic biodiversity in general and mahseers in particular. Integration of CPF, TPB, and social value analysis uncovered a broad stakeholder consensus on the decline of mahseer populations and the urgency of implementing conservation measures in Karnataka, Assam, and Uttarakhand. The stakeholders also urged to mitigate widely acknowledged threats faced by the natural mahseer populations such as dam construction, siltation, riverbed mining, and the invasion by alien fishes. It is well understood, bringing broad consensus among stakeholders is challenging but essential for any conservation interventions (Reed 2008). Identifying points of agreement alongside the state-specific issues of divergence, offers promising opportunities to formulate long- and short-term conservation goals at regional, state, and national levels as well the steps for achieving them (Reed 2018; Reed et al. 2014). Policymakers and aquatic ecosystem managers should therefore work towards translating this shared understanding present amongst diverse stakeholders into conservation actions. At the same time, topics where multistakeholder agreement emerged only in selected states - taxonomic status, species endemism and research leadership (Karnataka), jurisdictional overlaps (Karnataka and Assam), prioritisation of livelihood-oriented aquaculture (Assam), and penalties for illegal fishing (Uttarakhand) demand further research keeping the socio-cultural uniqueness of these regions in the background. Mapping such factors at micro-levels and integrating them into interventions can strengthen trust, support, and acceptance by the parties involved in the mahseer conservation, improve cooperation between them and reduce negotiation barriers (Pretty and Smith 2004; Brooks et al. 2013; Reed et al. 2014; Beever et al. 2019; Meierová 2020; Doley and Barman 2023). Moreover, this understanding can also help address inter- and intra-stakeholder conflict, a major impediment to conservation success (Pretty and Smith 2004; Reed et al 2014) and incorporate area-specific considerations into conservation strategies.

Although mahseer is a prized food fish and a popular recreational target across the focal states, stakeholders expressed strong social values, moral responsibility, and perceived behavioural control (PBC) toward protecting them. In Karnataka, multi-generational anglers and local communities displayed cultural and religious reverence for mahseers, and in Assam, conservation values were grounded in cultural and tribal identity and the long-standing fishing traditions. Meanwhile in Uttarakhand, mahseer held layered meanings - heritage symbol (State Fish), renowned sport fish, food resource, and a sacred entity (in a few regions). In spite of the fact that India has undergone colonial rule for centuries and here the conservation mindset and policies still bear the influence of the Western utilitarian views (Rangarajan 1999; Guha 2008) and human superiority over nature (Anderson and Perrin 2018; Fiasco and Massarella 2022), stakeholders including tribals, local communities and recreational anglers viewed fish, wildlife and nature as equal partners of humans, and kept a distance from the anthropocentric and exploitative concepts (Jyrwa et al. 2020; Iqbal et al. 2025). A large body of literature is available to demonstrate that values embedded in indigenous worldviews can unite diverse actors through knowledge systems rooted in cultural norms, traditions, and sustainable practices (Dhee et al. 2019; Jolly and Stronza 2025; Iqbal et al. 2025) and guide integrated socio-ecological solutions for coexistence (Tsing 2012; Ingold 2013; Haraway 2018; Ogilvie et al. 2018; Jolly et al. 2022; Shawon et al. 2025; Iqbal et al. 2025). However, even when attitudes, subjective norms, and PBC are strong, conservation behaviour remains limited when Actual Behavioural Control (ABC) is constrained (Ullah et al. 2021; Karimi and Ataei 2022), and fostering meaningful pro-conservation behaviour requires addressing decision-making power, adequate incorporation of stakeholder inputs, and structural constraints encountered by all parties involved, particularly the ground-level actors. As revealed by the present study, government officials across the focal states displayed lower ABC due to heavy workload, time limitation, and competing administrative priorities. Addressing these constraints through capacity building and establishing dedicated mahseer conservation teams actively collaborating with anglers, NGOs, and community leaders, could reduce the gap existing between PBC and ABC.

Despite these strengths, the mahseer conservation ecosystems in all three states studied were not free of shortcomings or conflicts, reminding that consensus on ecological problems did not always translate into agreement on solutions (Coglianese 1999). Misaligned values, ambiguity over jurisdictions, political interference etc. frequently triggered inter-stakeholder conflicts. In Assam, although cultural and tribal identity-based values supported mahseer conservation, inter-stakeholder frictions emerged often when livelihood security and socio-economic priorities clashed with conservation goals. In Uttarakhand, promotion of angling-based ecotourism and the establishment of temple-based fish sanctuaries (Dandekar 2013; Baruah et al. 2022) worked positively for mahseers, while heavy penalties for fishing in the lakes and rivers advised by The Uttarakhand Fisheries Act, 2003 (Uttarakhand Act No. 2 of 2003) was a reason for fisherfolks and locals not cooperating and frequently involving in the disputes with the authorities. In Karnataka, community associations and collaborative efforts between recreational anglers, researchers, the fisheries and forest Departments, and local communities have had a positive influence on mahseer conservation. However, conflicts have emerged due to differing views on mahseer endemicity and jurisdictional overlaps between the forest and fisheries Departments, resulting in divergent opinions and management challenges. Our results reiterate the need to balance livelihood options and community rights with the conservation efforts to ensure genuine stakeholder participation fostering long-term stewardship and sustainability (Newing et al. 2024). According to Reed (2008) conservation plans must consider entire riverine ecosystems, aligning species-level interventions with community well-being to achieve this goal. When conservation is perceived as externally imposed or culturally misaligned, compliance to the species management strategies and policies becomes superficial and short-lived (Pimid et al. 2022). Hence in Assam, a zoning approach allocating certain *beel* for aquaculture while reserving others for conservation breeding can accommodate both livelihood and mahseer protection. Recognising traditional subsistence fishing rights in National Park buffer zones in Assam and in the lakes and rivers of Uttarakhand, while maintaining bans on destructive methods and strengthening anti-poaching measures, could help reconcile competing interests. Reintroducing regulated angling-based ecotourism in Assam and establishing guidelines on permissible gear and catch limits, along with licensing systems for subsistence fishing, may further support this goal. Formally recognising researchers and recreational anglers as intermediary stakeholders can help mitigate jurisdictional conflicts between the fisheries and forest departments. Furthermore, institutionalising collaborative research and structured knowledge-sharing mechanisms among all stakeholders can address existing non-consensus on mahseer species endemicity in Karnataka and support more coherent conservation governance.

Collaboration (and communication) was another critical theme with notable interstate variation generated. Karnataka demonstrated robust micro-level partnerships among anglers, government departments, and local communities (Pinder and Raghavan 2013; Roy and Sreenivasan 2025), supported by breeding and ranching programmes, regulated angling, ghillie-led river patrolling, and fish sanctuaries declared under The Karnataka Inland Fisheries (Conservation, Development and Regulation) Act, 1996 (Karnataka Act 27 of 2003). In contrast, collaborations in Assam and Uttarakhand were limited and often ineffective due to poor interdepartmental coordination, political interference, bureaucratic challenges, exclusion of key actors from decision-making, and inconsistent involvement of local communities and NGOs. Such a fragmentation could undermine the trust and hinder knowledge exchange between stakeholders leading to the collapse of the conservation intervention as observed in the cases of Nameri Eco-camp and hatchery shutting down and the low impact of ‘Mahseer Watching’ in Assam. However, many success stories demonstrating the strength of active collaboration in promoting conservation are available from one of the focal states Karnataka, and the neighbouring states of Assam and Uttarakhand. Karnataka has successfully implemented regulated ecotourism and community monitoring of rivers in many areas of mahseers conservation importance generating benefits for local communities and helping mitigation of human–human conflict (Pinder and Raghavan 2013; Sreenivasan et al. 2021). Similarly in Meghalaya, a state located in the North Eastern India government-community partnership has led to the establishment of 54 fish sanctuaries since 2012 under the Meghalaya State Aquaculture Mission (MASM). Along with playing a crucial role in conserving the endangered golden mahseer (*T. putitora*) and near threatened chocolate mahseer (*N. hexagonolepis*) these community-led sanctuaries are also contributing to the protection of local rivers. With clear waters and thriving mahseer populations, these sanctuaries attract tourists thereby generating income for local communities efficiently monitoring and managing these water bodies (Dash et al. 2020; Rahman 2021; Jumani et al. 2023). Another collaborative initiative for fish conservation worth mentioning at this point is the ‘*Matsya Mitras’* (Friends of Fishes) launched by the Jharkhand State Government in 2007. Here, volunteers selected from the community and remunerated by the government work as a bridge between fisheries department, local fishing communities and NGOs. These volunteers disseminate new knowledge and techniques of aquaculture with fishermen and villagers interested in fish farming, document village-level water resources and thereby significantly enhance fish production and strengthen aquaculture development in the state (NITI Aayog n.d.; Shweta 2024). *Matsya Mitra* with local adaptation focusing mahseer could be useful for other states as well. Finally, validating roles of non-state actors reinforcing relational and moral values and environmental stewardship (Chan et al. 2016; Ewane 2024; Wato et al. 2025), and recognising and rewarding NGOs and community groups engaged in pro-conservation activities, through incentives, public visibility, or media coverage, can also strengthen stakeholder participation and accountability (Kipkeu et al. 2014; Rode et al. 2015; Ewane 2024).

Although effective communication between stakeholders is the backbone of the consensus, collaboration, and maintaining or modifying social values carried by the mahseer for positive changes in the behaviours of the stakeholders, across states, we observed departmental silos, communication gaps and absence of a common platform for sharing the thoughts. Presence of an active communication system is also essential for reducing human-human conflict (Huang et al. 2020; Wu et al. 2020) and promoting conservation action (Bickford et al. 2012). Furthermore, transparent communication with and between government departments is necessary to support informed decision-making and framing effective policies. Conservation Leadership Programme (CLP) in Baagi Village, Nayar River Valley in Uttarakhand (Mahseer School project, Dewan et al. 2022), stands out as an exception in this context. Through development of ‘Mahseer Schools’ and fostering an educational culture and curriculum that connects children to the fishes and nurture conservation ethic, this initiative have shaped pathways for the long-term restoration of golden mahseer (*T. putitora*) in the Ganga-Nayar river system (Dewan et al. 2022). In this state, many NGOs and government departments have employed creative communication strategies such as street plays, local radio broadcasts, newspaper articles, and wall graffiti to spread mahseer conservation messages. Expanding these approaches across states, together with organising regular inter-stakeholder meetings, could help counter existing communication gaps and to build trust and conservation literacy (Schultz and McGinn 2013) required for supporting mahseer and other indigenous fishes. A recent study exploring the mental models of multiple stakeholders related to mahseer conservation in Assam and Uttarakhand also highlighted the need for improved communication to facilitate inter-stakeholder collaboration, enhance efficacy of hatchery and conservation breeding operations, resolve human-human conflict, reduce illegal destructive fishing, and promote social values (Das and Binoy 2025b). In this context it is also important to remember the Targets 21 and 22 (Section H) and Section K of Kunming-Montreal Global Biodiversity Framework (CBD 2022) that respectively emphasises on participatory and inclusive biodiversity governance and recognises the importance of communication strategies that go beyond awareness to foster behaviour change through multiple-stakeholder engagement and effective messaging (CBD 2022). Therefore each state should consider forming mahseer and river conservation councils with units at different administrative levels and involving members from government departments, researchers, anglers, NGOs, and community representatives to improve coordination, resolve jurisdictional overlaps, and promote bottom-up decision-making. These organisations could also be developed as an inclusive platform for the stakeholders to exchange scientific information, discuss failures, and reconcile differences (Bickford et al. 2012). Such forums can cultivate and inculcate moral responsibility, promote Wildlife Value Orientations (WVOs) of mutualism and respect (Fulton et al. 1996; Gomez et al. 2022) amongst the stakeholders, along with helping fisheries department to communicate concerns about illegal fishing and local communities to voice livelihood-related challenges and bureaucratic obstacles they face. Furthermore, researchers should expand their circle of communication beyond academic publishing and share their findings and knowledge (Bagla and Binoy 2017) on mahseers with the public through social and traditional media in vernacular languages to improve mahseer and aquatic ecosystem conservation literacy. Our study also highlighted the importance of projecting mahseer’s religious and cultural significance while developing communication programmes and the importance of integrating local/tribal identity with such attempts (Clements et al. 2009; Bickford et al. 2012). Societal norms possess the power to override clear scientific evidence and hence a two-way engagement reflecting local socio-ecological realities and converging traditional knowledge with scientific insights is essential to foster shared ownership and more effective mahseer conservation (St. Clair 2003; Kahan 2010).

By connecting the structural steps of planning (Conservation Planning Framework, CPF) with social and psychological drivers of behaviour at the level of individuals (Theory of Planned Behaviour, TPB) and the cultural and social values with the potential to influence them (Chan et al. 2016; Marchini et al. 2019; Pimid et al. 2022) our study revealed that an approach integrating these frameworks can offer a holistic picture of the situation for designing and implementing conservation action plans. While CPF outlines the stages of assessment, decision-making, and implementation, it alone cannot capture stakeholders’ attitudes, subjective norms, or perceived behavioural control (TPB), which are critical for understanding why individuals or communities behave in a particular way. Addition of the details of social values, particularly relational values that reinforces moral responsibility and pro-conservation attitudes and hence the motivation for cooperation, and instrumental values shaping willingness to follow the changes advised by the conservation managers when livelihood and other benefits are at stake makes our framework vivid and wholesome. For instance, in Karnataka, Assam and Uttarakhand, while governance and regulatory clarity were the central drivers (CPF), their effectiveness depended heavily on attitudes and perceived and actual control of behaviours (TPB) of the stakeholders, as well as the alignment conservation plans had with their livelihood, cultural identity (social value), moral responsibility and stewardship (subjective norms, TPB). Furthermore, even when intentions were positive, low actual behavioural control (ABC - TPB) constrained many stakeholders from active participation in the conservation interventions resulting in suboptimal outcomes. Although the present study offers valuable insights into the human dimensions of mahseer conservation across states through a novel, integrated approach, it is not without limitations. Since current study is the first of its kind to integrate CPF, TPB and social value, more empirical and modelling studies should be conducted keeping different socio-ecosystems in the focus in order to strengthen the validity, and applicability of this framework. Increasing the number of stakeholders from each group studied and conducting exploration to make sure that views of all stakeholder groups are equally represented is also essential. Furthermore, possibility of the confirmation bias (Yeazitzis et al. 2023) that may arise due to unequal power relationships existing amongst the stakeholders and the political situations present in the focal communities also need to be managed to avoid reaching wrong conclusions. In spite of the limitations, our study has demonstrated that bridging CPF, TPB and social value frameworks helps to identify the points where social value orientations support or hinder behavioural intentions and hence offer insights to design technically sound interventions that are socially acceptable and capable of catalysing sustainable behavioural change, ultimately enhancing the effectiveness of mahseer conservation.

### Stakeholder Recommendations

Stakeholders across all three states proposed multiple recommendations for strengthening mahseer conservation through participatory, science-driven, and culturally grounded approaches, reflecting both shared and state-specific priorities. These recommendations are pointers for policy makers, authorities, conservation managers, and local bodies of governance to develop effective policies and programmes for protecting mahseer with national, state and regional implications. Jurisdictional confusion, reducing work load of the government staff, involving NGOs and anglers in decision-making, improving interdepartmental coordination, institutionalisation of the community-based co-management, promotion of culturally rooted stewardship etc. emerged as the topics demanding immediate attention. Other suggestions included establishing green no-fishing zones or fish sanctuaries on the line of Mahseer Conservation Reserve at Badi Lake, Udaipur (Rajasthan High Court 2017), ensuring legal protection for mahseers, enhancing inter-stakeholder communication, promoting transparent, bottom-up decision-making and allowing regulated subsistent fishing right with clear limits and licensing to the local communities. Many advised inclusion of the temple-based sanctuaries, and community-supported no-fishing zones within the respective State Fisheries Acts to align conservation targets with local practices. Expanding sustainable catch-and-release angling based eco-tourism and aqua-tourism as livelihood incentives for the stakeholders dependent on mahseer, strengthening and modernising mahseer breeding and ranching programmes and undertaking detailed mahseer distribution studies were also counted by many stakeholders as a need of the hour. Many suggested that awareness campaigns should move beyond one-way messaging and communicators should adopt culturally resonant, locally grounded strategies such as street theatre/*nukkad natak*, traditional songs etc., to strengthen the societal salience of the mahseers. Support for reframing traditional community fishing festivals as heritage events and transforming them into an opportunity to promote local-identify dependent mahseer conservation was also observed.

Recommendations received specific for each focal state were as follows: stakeholders from Karnataka highlighted the need for expanding citizen science and community-based initiatives, legitimising recreational angling as a conservation tool, strengthening training for forest guards, monitoring and managing the IAF and promoting collaborations between government departments, NGOs and researchers. They also emphasised declaring humpback mahseer (*T. remadevii*) the state fish and developing captive breeding programme for ensuring the seed availability of this endangered species, conducting genetic studies of different mahseer species present in the state, developing transitional pond ecosystems, and integrating mahseer’s cultural significance into conservation programmes. Conducting outreach through street plays and social media, issuing angling licences, allocating river beats to NGOs, and forming Indian researchers-led mahseer conservation groups were the suggestions recorded from Uttarakhand. Many stakeholders from this state also advised discouraging proposals for including mahseer under the Wildlife Protection Act, 1972, to safeguard livelihoods of locals and balancing strict penalties to encourage positive conservation behaviour. Mitigating political and bureaucratic challenges, empowering communities with fishing rights in buffer zones, licensing and training anglers for mahseer habitat monitoring roles, reframing community fishing events as heritage activities, and linking conservation with livelihood opportunities were the recommendations from Assam. All these suggestions reiterate that mahseer conservation policies should prioritise community-based and culturally grounded habitat management and discuss more rigorously the possibilities of integrating region specific aspects of freshwater ecosystem and aquatic biodiversity into the existing fisheries (Brooks et al. 2013; Oldekop et al. 2016) and wildlife protection Acts.

## 5. Conclusion

Human dimensions of successful conservation, especially those of freshwater ecosystems and the diversity it harbours, are deeply rooted in managing inter-stakeholder relationships through coordinated actions, inclusive communication, community empowerment, equitable resource allocation, and properly attending dynamics of the economical and political relationships present in the socio-ecosystems. Studying such complex scenarios demand an interdisciplinary approach integrating knowledge from the ecological, societal, cultural and individual dimensions. The present study was an attempt on this line to integrate the stages of the Conservation Planning Framework (CPF) with the Theory of Planned Behaviour (TPB) and social values to illustrate the conservation of charismatic megafishes, mahseer in three culturally and geographically different states of India. Along with highlighting the points of consensus our study threw light also on the misalignment of values and the topics on which inter and intra-stakeholder disagreement was conspicuous. Our analysis pointed out, addressing challenges of mahseer conservation requires inclusive, bottom-up decision-making and culturally embedded communication strategies that acknowledge moral and relational value dimensions of human-mahseer relationships. Furthermore, strengthening perceived behavioural control (PBC) and actual behavioural control (ABC) are also important as positive attitudes and norms alone are insufficient if actors lack autonomy, resources, or authority. Our findings demonstrated that integrating social values and individualistic behavioural dimensions into CPF can provide insights to foster human-mahseer coexistence, counter possible risks of biological and societal extinction of this iconic species, and to develop a national mahseer conservation policy with the scope to include region-specific demands and involve other socio-culturally and ecologically significant species across India and beyond.

## Acknowledgments

Prantik acknowledges the Human Research Development Group (HRDG) - Council of Scientific and Industrial Research (CSIR), New Delhi, India (09/1320(0001)/2020-EMR-I) for providing the research fellowship and the annual contingency grant for field-work related expenses. He extends his sincere appreciation and gratitude to all the respondents who took part in the interviews and FGDs across districts in all three study states. He also expresses his gratitude to Mrs. Bubu Sarkar Das, Mr. Subhashis Das, Dr. Monideepa Mitra and Mr. Mahendra Singh Rawat for their assistance in facilitating stakeholder contacts and data collection in Uttarakhand; to Ms. Ankita Sharma, Dr. Achyut Malakar, Mr. Abhijit Konwar and Mr. Dhireswar Sarma in Assam for additional assistance with Assamese translations; and to Mr. Balaraj Hiremath in Karnataka for additional assistance with Kannada translations.

## Author Contributions

**Prantik Das:** Conceptualisation; methodology; investigation; data curation; formal analysis; visualisation; funding acquisition; writing - original draft; writing - review and editing. **V. V. Binoy:** Conceptualisation; methodology; visualisation; writing - review and editing; supervision.

## Informed Consent Statement

All interviews and FGDs were undertaken in compliance with the institutional ethical guidelines for conducting non-invasive research involving human participants, as approved by the Institutional Ethics Committees (IEC) of The University of Trans-Disciplinary Health Sciences and Technology (TDU; Protocol Number: TDU/IEC/16E/2025/PR72) and the National Institute of Advanced Studies (NIAS; Letter Number: NIAS-EC-11/03/2022). In addition, the field study carried out in the Kosi Forest Range, Ramnagar, Nainital was undertaken after obtaining formal permission from the Divisional Forest Officer (DFO), Ramnagar Forest Division, Nainital, Uttarakhand (Letter Number: 2873/61), in accordance with the rules of the forest department for this specific region. Prior to the participation, all respondents were made fully aware about the objectives of the study, and it was emphasised that their participation was completely voluntary, with the option to withdraw from interviews or FGDs at any stage. The data collection through audio-recordings, note-taking and photography was carried out only after obtaining oral/audio-recorded or written informed consent. The participants were also assured that their names and other personal details that could be used to identify them, would remain confidential and anonymous to protect their identity and privacy.

## Data Availability Statement

In order to maintain the privacy of the respondents, the datasets have not been made publicly available, as they contain information that could compromise research participant consent. Accordingly, the data used in this study remain confidential.

## Supplementary Materials (SM)

**SM 1.**
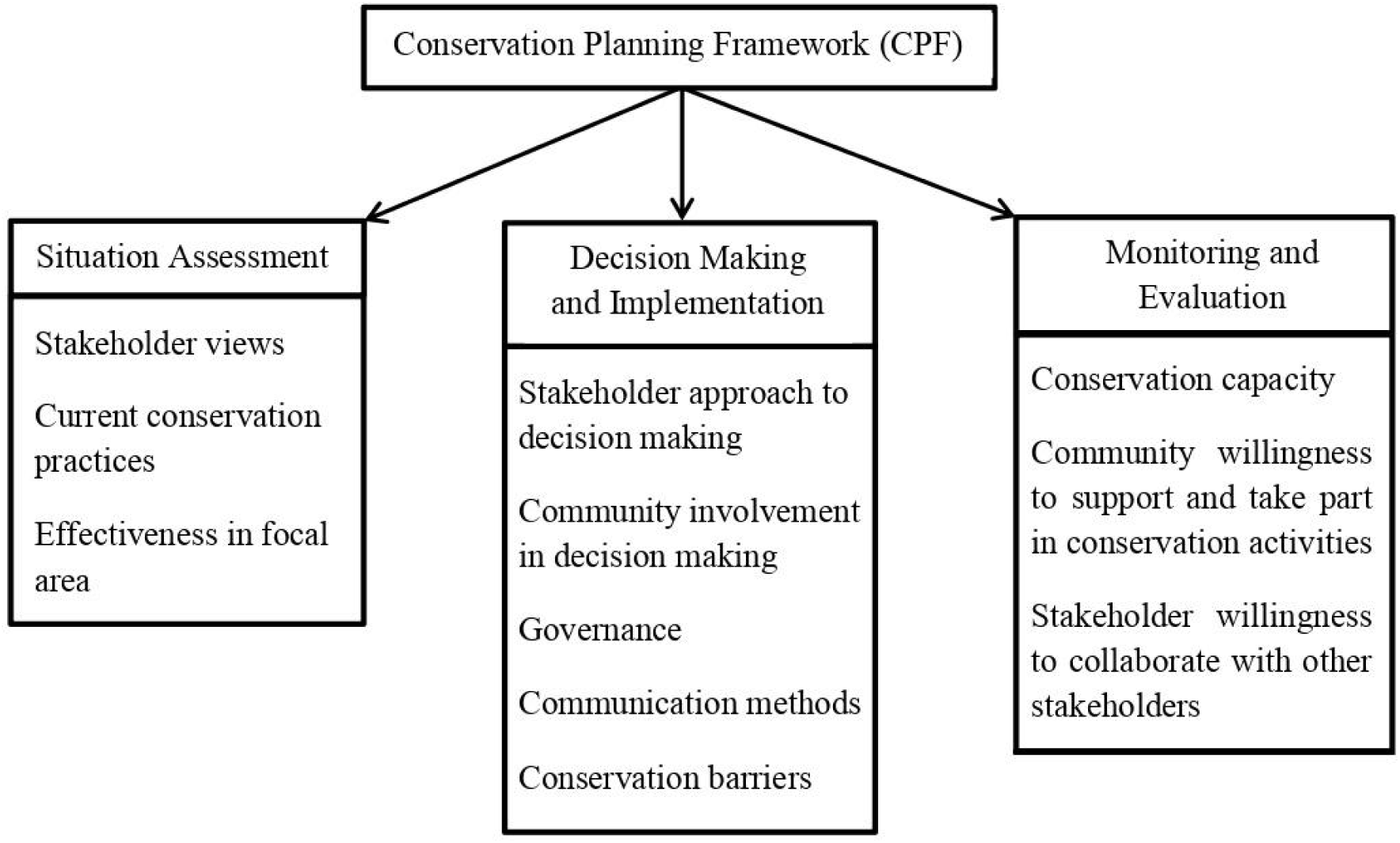
Components of Conservation Planning Framework (CPF; Marchini et al. 2019)

**SM 2.**
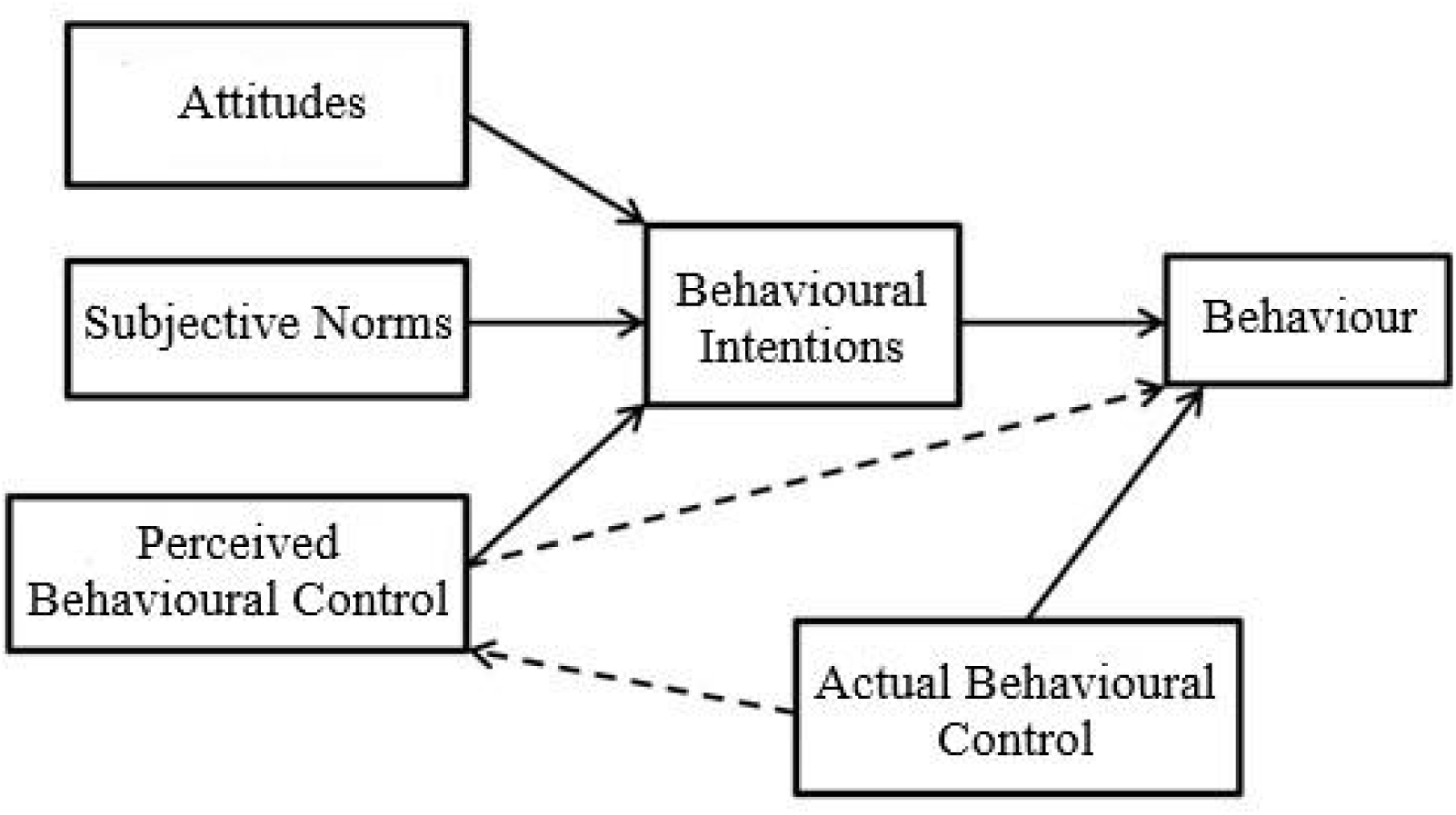
Components of Theory of Planned Behaviour (TPB; Ajzen 1985)

SM 3. Stakeholder Interview/FGD Questions:

## Section 1: Conservation Planning Framework (CPF)

1. Are you aware of different mahseer species found in your state? What are they called in the local language?
2. (If unaware) Based on historical records, they were present in this region in the past. Is it correct that they occurred here earlier but are not found currently?
3. Do you think the number of mahseer (species name changed according to the state where the interview is being conducted) is declining in the rivers in your locality?
4. What threats do you think mahseers face in your locality?
5. Do people practice any destructive fishing methods here? What are those?
6. Do you think that traditional community fishing is impacting the local fish population?.
7. Do you know what the non-local Invasive Alien Fishes (IAFs) are? Do you think that IAFs have any effect on the wild mahseer populations in your locality?
8. Should there be stocking of IAF in rivers where mahseer population thrives for the economic benefit? Why/Why not?
9. What are your views on recreational angling of mahseer? Is it beneficial or harmful?
10. If beneficial, then how have they helped?
11. Do you believe that anglers and the mahseer can thrive together sustainably? How in your opinion can it be sustainable for both the fish and the anglers?
12. Are there any official guidelines or educational materials from the government regarding proper angling techniques? What is your opinion on that?
13. Are you aware about the IUCN status of the mahseer species?
14. Do you think that there is a need to conserve the mahseer species? If so, any opinions on which mahseer species?
15. What are the existing mahseer population management programmes that are being undertaken by the Department (Fisheries and Forest) in the state? Are they separate for different mahseer species?
16. Do you think these have been effective in leading to a positive outcome? Do you feel any need for change in these programmes?
17. Do you think interventions are required for the conservation of the mahseer population? What interventions can be done to bolster the stock in the wild?
18. Is there any particular species of mahseer in your opinion that needs more conservation efforts? Why?
19. Mahseer fish species are not included as ‘wild animals’ under the schedule of the Wildlife Protection Act of 1972 in Section 2(A). What is your opinion on that?
20. Is subsistence fishing allowed for the locals within the National Park/Wildlife Sanctuary/protected area like mahseer fish sanctuary?
21. If yes, is there a licensing system in place?
22. Have the local community members or other stakeholders (such as researchers, anglers) been involved in the decision making process related to mahseer conservation plans and strategies?
23. Does the stakeholder hold open discussions with the local community members or involve them in awareness activities regarding mahseer conservation?
24. Does the stakeholder work in association with the other stakeholders such as research institutions, recreational anglers, NGOs, government departments with respect to the mahseer conservation programs?
25. Are there sufficient resources (human and monetary) for dedicated mahseer conservation?
26. What are the conservation communication and awareness activities being undertaken by the stakeholders? Do the locals support these activities?
27. How important do you think conservation communication is for mahseer conservation activities to be successful?
28. Do you think social media can be used as a tool for communication?
29. What is your opinion on the creation of fish sanctuaries or no fishing green zones?

## Section 2: Theory of Planned Behaviour (TPB)

1. In your view, do the stakeholders possess adequate knowledge and information to effectively undertake mahseer conservation efforts on their own? Is that knowledge shared with other stakeholders?
2. Is there any kind of social pressure from your colleagues/government/family members/head of the community to involve in mahseer conservation related activities?
3. In your opinion, does the existing official workload limit the capacity of the staff to focus specifically on mahseer conservation efforts? (For Government stakeholders)
4. What does mahseers mean to you, do you either see it from the point of view of conservation or a way for economic benefits or a combination of both?
5. Are you involved with conservation activities? If not, how willing would you be to get involved in conservation initiatives in your area?
6. Do you collaborate with other stakeholders in such activities?
7. What personal needs, interests, or motivations might drive you to get involved in these kinds of initiatives?
8. What would you say, in your opinion, are the barriers to mahseer conservation?

## Section 3: Social Values

1. What is your opinion on the religious and cultural significance of mahseer species (in the area where the interview is being held)?
2. Are mahseers a part of any cultural or mythological stories?
3. In your view, should such cultural values be integrated into the design of region-specific mahseer conservation plans, such as temple fish sanctuaries? How effective would those be?
4. What kind of socio-economic benefits have mahseer provided to the community over generations?
5. Why do you think that mahseers should be protected and conserved?
6. What kind of non-tangible, non-monetary benefits, in your opinion, do mahseers provide to you and the society?
7. How do you think stakeholders view mahseer conservation — as a personal commitment to nature and biodiversity conservation, a livelihood opportunity, a reflection of cultural or religious identity, a job responsibility (professional obligation), or a combination of these?
8. Are there any additional recommendations you would like to suggest to enhance the effectiveness and efficiency of the existing mahseer conservation protocols and strategies?

### Additional questions for recreational anglers

1. How long have you been angling? Which river systems?
2. During which months do you angle? Is there a closed season in the rivers for angling?
3. Do you often catch IAF as by-catches when angling mahseer?
4. Do you change your angling gear every angling season? Should it be followed by all anglers?
5. What are your thoughts on handling of fish, time of fish in air, proper hooks and lines (different aspects of angling)?
6. In 2011, a legislation to ban Catch & Release (C&R) from protected areas was passed. Did you know about that? What is your opinion on that?
7. Do you need to take a license to angler in the rivers of the particular state? Who issues the license?
8. Do you practise Catch & Release (C&R)? Do you think C&R is an ethical tool for angling? Do you think the percentage of anglers practising C&R is high or low?
9. What is your opinion on the anglers’ beliefs that they feel majorly responsible towards the fisheries?
10. Do you think that the anglers should feel that they have a duty towards the mahseer conservation or do you think that they should see angling from the point of being just another recreational sport?
11. Do you think there is a potential to involve/motivate more people in angling through your social media posts?
12. Any negative impact of social media angling posts that you have faced first hand? Has it affected your angling practice or mahseer populations?

### Additional questions for Fisheries Department staff

1. Certain stretches of rivers have been declared as a Fish Sanctuary? What is the length of the Sanctuary? Which year was it declared as a Sanctuary? By whom? What kinds of activities are banned along the stretch?
2. Certain stretches of the rivers have been leased? Who is eligible to apply for this lease? What is the length of the lease stretch? When and why was it leased? Are there any conservation activities that are being done by the Fisheries Dept. along that stretch leased?
3. How is this stretch leased different from the stretch declared as a Fish Sanctuary? Do they overlap?
4. What are the activities done in the hatchery? (breeding, rearing, selling, distribution without cost)
5. Which mahseer species are being bred in the hatchery? (If only one species) Why are the other species not bred? Why has the hatchery only focused on one specific mahseer species?
6. Were there efforts to breed others? Any future plans to breed them?
7. Do the officials train the fish farmers regarding maheer or only for IMC aquaculture?
8. Do you believe that housing the mahseer and the IAF fingerlings together in the hatchery would help in better survival in the wild?
9. Do you provide fingerlings to local people? For what purpose?
10. Do you think the present hatchery/breeding protocol is enough for conservation? Are there any gaps in the system?
11. Do you think the goal should be to increase the already massive productivity or to create a self-sustaining population in the wild?
12. Has the hatchery protocol changed over the years? (If yes) At what level of breeding did it change? (If not) Do you feel any need for changes in that?

### Additional questions for Forest Department staff

1. Are there any targeted mahseer conservation plans or they are protected as part of the forest and biodiversity?
2. Do you conduct a creel census to determine the number of individuals?
3. Are the forest guards consulted during discussions with the ranger or DFOs?
4. Do the forest guards face any kind of conflict from the local residents of the buffer zone?
5. Are the buffer zone residents invited for discussions over decision making?
6. Are the buffer zone residents permitted to subsistence fish with rod without any destructive methods? Were the rules the same before or have they changed?
7. Do outsiders come to perform community fishing in the river systems of the protected areas?
8. What do you think is the ideal way to interact with the buffer zone residents demanding the fishing rights?
9. Is angling allowed as part of eco-tourism activities within the buffer zone? Have any of the rules changed?
10. Do you provide any licenses to local fishermen for fishing in the buffer zone? If yes, how many licenses are provided? Who are they provided to?
11. Is there any closed season for fishing?
12. What are punishments for illegal destructive fishing or poaching?

### Additional questions for researchers

1. Are you invited to discussions over mahseer decision making?
2. Does your research have a component of intervention as well?
3. Have you conducted any intervention? How effective have those been?
4. What are the scientific barriers to mahseer in breeding?
5. Have you been consulted during breeding or reintroduction or stocking programmes or research work?
6. Have you faced any conflict with any stakeholder during your research work which included field-work?

### Additional questions for fishermen

1. What kind of fishes do you generally catch?
2. Do you need a license to fish in certain stretches of the river? What are these stretches? Who issues the license?
3. Do you catch mahseer? If yes, do you land them frequently?
4. Have the number of mahseer catches changed over time? What about the IAFs?
5. What do you do if you land a mahseer? Are they taken for consumption purposes?
6. Do you also farm or breed fishes for livelihood purposes?
7. Are you consulted by the fisheries department officials on what kind of fish to be bred?
8. Do discussions happen with the fisheries department?
9. How important is involvement in conservation activities for you along with the livelihood based fishing and aquaculture?
10. Have you seen destructive fishing being undertaken? Have they declined or increased over time?
11. Along with training regarding IMC breeding and aquaculture, are you provided with training or education regarding conservation of endemic fish species?
12. In case you are asked to involve yourself in conservation activities, will you be willing to do so? Are there sufficient people for undertaking such activities?

### Additional questions for local community members involved in community fishing events

1. What is a community fishing event? When are they organised during the year?
2. Which community participates in this event? How is it conducted?
3. Which water bodies are they conducted in, rivers or lakes?
4. How important is it as a part of heritage and culture?
5. There have been reports of impact on the fishes due to these activities. What is your opinion on that?
6. Have there been any conflicts with any officials or NGOs regarding community fishing?
7. Do you get mahseer species in your catch during the event?
8. Have the number of fish caught declined over the years? Do you get IAFs in your catch?
9. Has this resulted in greater tourism and acted as a source of revenue for the local community?
10. Do you get poachers or people engaging in destructive fishing methods during this festival? What do you do in such instances?
11. Who are the decision makers regarding community fishing?
12. Are the local elected members or government officials involved in the decision making?
13. Do you interact with local NGOs and researchers?
14. Have you been involved in spreading awareness for practicing sustainable community fishing?
15. Does the media focus on these events? If yes, what dimension of the event do they mostly focus on?
16. Have you seen any changes in how the event is conducted over the years? Are those positive changes or negative?
17. In your opinion, how can the community fishing events be modified or moulded to promote sustainability?
18. At the same time, how can they be modified so that the respect towards the traditions and culture of the region stays intact?

